# Otenabant is a Selective Antagonist of Human PIEZO1

**DOI:** 10.64898/2026.04.28.720360

**Authors:** Vincent Jaquet, Reetta Penttinen, Gorane Rodríguez-Urquirizar, Cyril Castebou, Yves Cambet, Nicolas Rosa, Mathilde Bourdin, Bita Asghariastanehei, Franciele de Lima, Marie Martin, Elie Nader, Selma A. Serra, José M. Fernández-Fernández, Nicoletta Murciano, Philippe Connes, Stéphane Egée, Hélène Guizouarn, Lars Kaestner, Niels Fertig, Maria Giustina Rotordam, Nicolas Demaurex

## Abstract

**Background and purpose:** PIEZO1 mechanosensitive cation channels translate mechanical cues into intracellular Ca^2+^ and Na^+^ elevations, enabling cells to respond to physical alterations in their environment. PIEZO1 contributes to red blood cells (RBC) volume homeostasis and gain-of-function PIEZO1 mutations cause hereditary xerocytosis (HX), a rare mostly compensated hemolytic anemia, and aberrant channel activation exacerbates sickling and vascular dysfunction in sickle cell disease. Despite strong genetic and physiological evidence supporting PIEZO1 as a therapeutic target, potent and selective inhibitors are limited, and existing compounds show modest specificity or poorly explored mechanisms. Improved pharmacological tools are needed.

**Experimental approach:** We conducted a high-throughput screen of FDA-approved drugs to identify PIEZO1 inhibitors. Compounds were tested at concentrations of 10 µM in a monocytic cell line, using intracellular Ca^2+^ elevations evoked by the PIEZO1 agonist Yoda1 as read-out. The inhibitory activity of the best hit was validated and compared to existing PIEZO1 inhibitors using electrophysiological analysis, orthogonal PIEZO1-dependent assays across cell lines and human RBCs. As functional proof, we investigated the impact of three PIEZO1 inhibitors on RBC deformability by ektacytometry, after Yoda1 pre-stimulation.

**Key results:** This screen identified Otenabant, a selective Cannabinoid Receptor Type 1 (CB1) antagonist, as a potent PIEZO1 inhibitor. Otenabant dose-dependently inhibited Ca^2+^ elevations mediated by endogenous or exogenously expressed human PIEZO1, but was ineffective against mouse Piezo1, revealing species-specific channel differences. Otenabant inhibited mechanosensitive currents elicited by shear stress in fibroblasts and by repeated poking in PIEZO1-expressing HEK-293 cells, altering the currents activation and inactivation kinetics, and prevented Yoda1-induced hyperpolarization in RBCs. Otenabant was able to reverse the negative impact of Yoda1 on RBC deformability.

**Conclusions and implications:** These findings demonstrate the utility of Yoda-based screening for discovering PIEZO1 antagonists and identify Otenabant as a promising chemical scaffold for developing selective PIEZO1 inhibitors with therapeutic potential.

## Introduction

Mechanotransduction is a fundamental biological process by which cells convert mechanical forces into biochemical signals. Through this process, mechanical stimuli activate intracellular signalling pathways that drive key adaptive responses. With the discovery of PIEZO1 and PIEZO2, researchers gained crucial insight into the mechanisms by which mechanical forces govern cellular function (Coste et al., 2010). Upon mechanical activation, such as poking, stretch or shear stress, PIEZO1 and PIEZO2 channels translate membrane tension into intracellular Ca^2+^ elevations, enabling cells to sense and respond to their physical environment.

PIEZO2 is predominantly expressed in Merkel cells and sensory neurons, where it mediates touch sensation and proprioception (Woo et al., 2014). In contrast, PIEZO1 exhibits widespread expression in non-sensory tissues, including red blood cells (RBCs), immune cells, and keratinocytes (Karlsson et al., 2021), where it regulates various mechanosensitive processes, such as contribution to RBC volume regulation (Kuck et al., 2025). PIEZO1 activation typically requires direct mechanical stimulation of the cell membrane, making functional studies technically demanding. Yoda1, a selective small-molecule agonist, allows pharmacological activation of PIEZO1 by shifting mechanical sensitivity, effectively mimicking mechanical force and facilitating scalable experimentation (Syeda et al., 2015).

Mounting evidence implicates PIEZO1 dysregulation in various pathological conditions characterized by enhanced Ca^2+^ influx. In RBCs, PIEZO1 contributes to cell volume and deformability regulation and gain-of-function variants cause hereditary xerocytosis (HX), a rare mostly compensated hemolytic anemia in which excessive PIEZO1-mediated Ca^2+^ entry triggers Gárdos channel activation and subsequent cellular dehydration (Zarychanski et al., 2012). PIEZO1 activation also exacerbates RBC sickling and adhesiveness in sickle cell disease, contributing to vascular occlusion and hemolysis (Nader et al., 2023) (Romero et al., 2025). Moreover, numerous studies have shown that conditional Piezo1 deletion in mouse specific tissues mitigates key pathologic features of several disease models with unmet medical needs, including idiopathic pulmonary fibrosis (Xu et al., 2025), and melanoma metastasis (Silvani et al., 2025) among others.

Despite growing interest in PIEZO1 as a therapeutic target, potent and specific inhibitors remain scarce. The peptide toxin GsMTx4, derived from a tarantula venom, is the most widely used PIEZO1 inhibitor but acts indirectly by modifying membrane tension rather than binding PIEZO1 directly (Gnanasambandam et al., 2017). Other inhibitors, such as Ruthenium Red and Gd^3+^, lack selectivity, while *Dooku1* antagonizes Yoda1 but not mechanically induced activation (Evans et al., 2018). Additional compounds, including Jatrorrhizine (Hong et al., 2023), Salvianolic acid B (Pan et al., 2022), SC-560 (JP2014084283 (A)), and Benzbromarone (Liang et al., 2024), have been reported to inhibit PIEZO1, but their mechanisms and pharmacological specificity remain incompletely characterized. Notably, Benzbromarone reverses echinocyte formation in HX patient-derived RBCs, validating its clinical potential (Liang et al., 2024).

The current lack of validated, selective PIEZO1 inhibitors highlights the need for new chemical tools to probe PIEZO1 biology and evaluate its therapeutic potential. In this study, we performed a high-throughput screen of a well-characterized library of FDA-approved drugs to identify compounds capable of inhibiting PIEZO1 activity, using Yoda1-induced Ca^2+^ influx as a functional readout. From this screen, we identified Otenabant, a selective cannabinoid receptor CB1 antagonist, as a potent PIEZO1 inhibitor. The inhibitory efficacy of Otenabant on PIEZO1-mediated Ca^2+^ entry was compared with that of the reference inhibitors Benzbromarone and SC-560. PIEZO1 inhibition by Otenabant was further validated through the patch-clamp technique and different orthogonal PIEZO1-dependent assays, including functional studies using human RBCs.

## Methods

### Reagents

CP-945598 HCl (Otenabant) (CAS 686347-12-6), Chem-space; benzbromarone (CAS 3562-84-3), MedChem Express; SC-560 (CAS 188817-13-2), AmBeed; Yoda 1 (CAS 448947-81-7), Sigma-Aldrich; Yoda 2, hellobio; Jedi2 (CAS 651005-90-2), Sigma-Aldrich; Jatrorrhizine (CAS 6681-15-8), Cayman; Salvianolic acid (CAS 121521-90-2), TCI; GsMTx4 (CAS 1209500-46-8), MedChem Express; Gadolinium (GdCl3) (CAS 13450-84-5), Sigma-Aldrich; Dooku1 (CAS 2253744-54-4), Sigma-Aldrich, Taranabant (CAS 701977-09-5), AM251 (CAS 83232-66-8), AM4113 (CAS 614726-85-1), MedChem Express; Thapsigargin (CAS 67526-95-8), Sigma-

Aldrich; L-012 (CAS 143556-24-5), IG instruments; Cal-520, AM, AAT Bioquest; Lipopolysaccharide (CAS 93572-42-0), Sigma-Aldrich; Doxyxycline (CAS 10592-13-9), Sigma-Aldrich; Penicillin/Streptomycin, Gibco; ATP, De Revvity.

### Cell culture

PLB-985 (RRID: CVCL_2162) were kindly provided by Prof. Karl-Heinz Krause, University of Geneva, Switzerland. PLB-985 cells were grown in suspension in 75 cm^2^ culture flasks in RPMI-1640 medium (GIBCO; #61870–010) containing 10% fetal bovine serum and penicillin/streptomycin (50 U/mL) at 5% CO_2_/95% air in a humidified atmosphere at 37 °C. HT1080 cells (RRID: CVCL_0317) were kindly provided by Dr Patrick Salmon, University of Geneva, Switzerland. They were grown in DMEM 4.5 g/mL containing 10% fetal bovine serum and penicillin/streptomycin (50 U/mL) at 5% CO_2_/95% air in a humidified atmosphere at 37 °C. Mouse embryonic fibroblasts (MEF) were kindly provided by Prof. Luca Scorrano, University of Padua, Italy and were grown in DMEM medium, 1 g glucose/mL GIBCO 22320 containing 10% fetal bovine serum and penicillin/streptomycin (50 U/mL) at 5% CO_2_/95% air in a humidified atmosphere at 37 °C. JawsII (RRID: CVCL_3727, ATCC® #CRL-11904™) were grown in suspension in 75 cm^2^ culture flasks in MEM Alpha medium (GIBCO), 4 mM L-glutamine GIBCO cat#25030, 1 mM sodium pyruvate SIGMA cat#58636 stock, 5 ng/mL mL murine GMCSF Immunotool cat#12343125 containing 10% fetal bovine serum and penicillin/streptomycin (50 U/mL) at 5% CO_2_/95% air in a humidified atmosphere at 37 °C. Adherent cells (MEF and HT1080) were passaged every 3–4 days using trypsin 0.25%/EDTA (GIBCO, #25200-056) at 25,000 cells per cm^2^. HEK T-REx™ 293 cells overexpressing either human PIEZO1 (hPIEZO1, (Rode et al., 2017)) or murine Piezo1 (mPiezo1, (Blythe et al., 2019)) were kindly provided by Prof. David J. Beech, University of Leeds, UK. For automated patch-clamp recordings cells were cultured and harvested according to Nanion’s standard protocols (Obergrussberger et al., 2014).

### Stable cell line generation

*Piezo1* knock-out cell lines were created by CRISP-Cas9. gRNA sequences targeting either human (GTG CAA GCA GTG TTA CGG CC) or mouse (AAT GAC CAG CGT CCG TCC AG) Piezo1 were cloned into a lentiGuide-Puro plasmid (Addgene #80263) using the Q5 mutagenesis kit (New England Biolabs) with the following primers: TGT TAC GGC CGT TTT AGA GCT AGA AAT AGC AAG (human fw), CTG CTT GCA CCG GTG TTT CGT CCT TTC C (human rv), GTC CGT CCA GGT TTT AGA GCT AGA AAT AGC AAG (mouse fw) and GCT GGT CAT TCG GTG TTT CGT CCT TTC (mouse rv). All constructs were verified by Sanger sequencing (Fasteris). For Cas9 expression, we used our previously generated piRFP720-Blasti-Cas9-FLAG plasmid (Carreras-Sureda et al., 2023).

For the generation of the Piezo1^KO^ PLB-985 cell line, cells were transiently transfected with the lentiGuide-Puro-Piezo1 and the piRFP720-Blasti-Cas9 plasmids using an Amaxa Cell Line Nucleofector (Lonza) following manufacturer’s instructions for HL-60 cells (Kit V, Program T-019). Four days after transfection, iRFP720-positive cells were sorted using a MoFlo Astrios cell sorter (Beckman Coulter). Efficient PIEZO1 depletion was validated by the absence of response to Yoda2 in single-cell Ca^2+^ imaging and by western blot.

For the generation of the PIEZO1KO HT1080 and MEF cell lines, we first generated cell lines stably expressing the Cas9. Cells were transduced with lentiviral particles produced in HEK293T cells (RRID: CVCL_0063) by co-transfection of the piRFP720-Blasti-Cas9, psPAX2 and pMD2.G plasmids. HEK293T transfection, lentiviral particle collection and transduction were performed as described in (Salmon, 2013). After one week of blasticidin selection, iRFP720-positive cells were sorted as described above. Stable Cas9 expression was confirmed by western blot. The Cas9-expressing cells were later transduced with lentiviral particles produced in HEK293T cells by co-transfection of the lentiGuide-Puro-Piezo1, psPAX2 and pMD2.G plasmids. After one week of puromycin selection, 96 single clones were isolated in a 96-well plate using a MoFlo Astrios cell sorter (Beckman Coulter). Following amplification, Ca^2+^ response to Yoda2 was measured for each clone using a FDSS/μCELL plate reader (Hamamatsu). Efficient PIEZO1 depletion in the non-responsive clones was further validated by single-cell Ca^2+^ imaging and by western blot.

The stable MEF-GenEPi cell line was generated by transposition of the GenEPi (human PIEZO1 with a low affinity GCaMP sensor (Yaganoglu et al., 2023), coding sequence along with a Tet-On promoter into MEF-Piezo1KO cells. First, we created a Gateway entry vector containing the GenEPi coding sequence: pENTR1a-GenEPi. We performed a PCR with the following primers fw: GCT CAA GCT TAT CGC ATG GAG C and rv: TAT GAT CTA GAG TCG CGG CCG on a pCMV-GenEPi plasmid (Addgene #140236) to amplify the GenEPi sequence, and another PCR with the following primers fw: ATT AGC GGC CGC TCG AGA TAT CTA GAC C and rv: CAT AAA GCT TGC AGC ATG GTG GCG ATC on a pENTR1a-tagRFP-KDEL (Addgene #114177) to isolate the pENTR1a backbone. The extremities of the two DNA products were cut with the restriction enzymes HindIII and Not1 (New England Biolabs) and ligated with the T4 DNA ligase (Thermo Fisher Scientific) according to manufacturer’s instructions. In parallel, we generated a transposable and tetracycline-inducible expression system based on the Sleeping Beauty transposon system (Kowarz et al., 2015) and on the polyswitch Tet-On vector (Giry-Laterriere et al., 2011) pCLX-pTF-DEST-EBR. We performed a PCR with the following primers fw: CAT AGG TAC CTC TAG GCG AGA CCC TGT C and rv: GAT AGC TAG CAA ACT ACC CCA AGC TGG C on a pSBbi-Pur plasmid (Addgene #60523) to amplify the Sleeping Beauty backbone without its constitutive promoter, and another PCR with the following primers fw: GCA TGG TAC CTT CGG TAC AAC TCC CAC and rv: ATA AGC TAG CAG CTC GAA TTC CAG GCG on the pCLX-pTF-DEST-EBR plasmid (gift from Dr. Patrick Salmon, …) to isolate the pTF-DEST-EBR inducible promoter system. The extremities of the two DNA products were cut with the restriction enzymes KpnI and NheI (New England Biolabs) and ligated with the T4 DNA ligase (Thermo Fisher Scientific) according to manufacturer’s instructions. As the newly generated pSbi-pTF-DEST-EBR plasmid contains a Gateway destination sequence, we performed a Gateway cloning reaction with the pENTR1a-GenEPi donor plasmid using the LR clonase II (Invitrogen) to generate the pSbi-pTF-GenEPI-EBR. All PCR were performed using the Herculase II Fusion DNA Polymerases (Agilent) according to manufacturer’s instructions. Primer annealing temperature was determined using IDT’s OligoAnalyzer™ primer analysis tool (https://eu.idtdna.com/pages/tools/oligoanalyzer). All constructs were verified by Sanger sequencing (Fasteris). Finally, MEF-Piezo1KO cells were co-transfected with the pSbi-pTF-GenEPI-EBR and pCMV(CAT)T7-SB100 (Addgene #34879) plasmids using Lipofectamine 2000 (Invitrogen) according to manufacturer’s instructions. The SB100X transposase triggered transposition of the pTF-GenEPI-EBR cassette into the MEF genome. After one week of puromycin selection, we obtained a tetracycline-inducible MEF-GenEPi stable cell line.

### Western blotting

Total protein extraction and western blots were performed as previously described (Rosa et al., 2022). PIEZO1, vinculin and HSP90 were detected using mouse monoclonal antibodies

(respectively: Invitrogen MA5-32876, Sigma-Aldrich V9264, and Santa Cruz Biotechnology sc-13119) and HRP-coupled anti-mouse antibodies (BD Pharmingen 554002, RRID: AB_395198).

### Ca^2+^ measurements

Ca^2+^ measurements were performed using the FDSS functional drug screening µCell kinetic plate reader. Cells were loaded (2 × 10^6^ cells/mL) with 4 μM Cal-520, AM in CA buffer (in mM: 140 NaCl, 5 KCl, 1 MgCl_2_, 20 HEPES, 2 CaCl_2_, 10 glucose, pH 7.4) for 45 min at room temperature in the dark, then cells were washed and resuspended in CA buffer at 10^6^ cells/mL PLB-985. Loaded cells (50,000 per well in 40 μL) were dispensed in 384-well microplates (corning #3770). Baseline fluorescence (Excitation 485 nm/emission 520 nm) was measured for 1 min every 500 ms. Compounds (5 µL diluted in CA buffer) at 50 µM were added on cells during acquisition. Fluorescence was recorded for 10 min before addition of 5 µL of Yoda1 (50 µM) in CA buffer and measured for additional 5 min.

### High-throughput screening

The library of 1971 of FDA-approved small molecules was purchased at APEXBIO. Compounds were stored in dimethylsulfoxide (DMSO) solution at 10 mM and kept at -20 °C.

The screen was performed in 384-well format in biological duplicates. Compounds were added in columns 3-22 of 384-well plates, while columns 1, 2, 23 and 24 were dedicated for controls. Hits were defined as less than 30% of positive control (Yoda1 alone).

### Reactive oxygens species (ROS) detection

PLB-985 cells (2 × 10^5^ cells/mL) were differentiated into granulocyte-like cells by addition of DMSO (1.25% final concentration) for 4–5 days. The assay was performed in a volume of 50 μL per well in white nonbinding surface 384-well plates (Corning). Differentiated PLB-985 cells (10^6^ cells/mL) were resuspended in HBSS buffer (GIBCO #14025050) supplemented with L-012 (50 µM final, Dojindo) and dispensed into the 384-well plate containing serial dilutions of compounds (1% DMSO) for 10 min. NOX2 was activated by addition of phorbol-12-myristate-13-acetate PMA (10 nM final). Luminescence was measured with a Spectramax Paradigm reader (Molecular Devices) for 1 h. To graph the data, area under the curve (AUC) was calculated.

### TNF ELISA

PLB-985 cells WT and PIEZO1 KO (2 × 10^5^ cells/mL) were differentiated into granulocyte-like cells by addition of DMSO (1.25% final concentration) for 4–5 days. Cells (50,000 cells/well in 100 µL RPMI) were dispensed in a 96 well plate, stimulated with LPS 1 µg/mL for 4 h and Yoda1 10 µM and Otenabant (OTB) and Benzbromarone (BBR) (10 µM) were added for 1 h. Supernatant was collected and 50 µL were used for TNF detection using a Human TNF alpha Uncoated ELISA kit from Invitrogen.

### Automated patch-clamp recordings

Automated patch-clamp recordings were conducted using the SyncroPatch 384 (Nanion Technologies GmbH, Munich, Germany) and NPC-384 multi-hole chips on HT1080 WT and P1KO cells (4-holes per well, 0.5-0.9 MΩ resistance) and on HEK T-REx™ 293 hPIEZO1 and mPiezo1 cells (8-holes per well, 0.6-1.2 MΩ resistance). Each well of the chip is connected to an individual amplifier channel, thereby enabling independent patch-clamp recording. The total current recorded from each well represents the sum of currents generated by single cells captured on the apertures. Recordings were performed at 26 °C (mPiezo1) or 30 °C (hPIEZO1 and HT1080 WT and P1^KO^) using the perforated-patch configuration with 8 µM (mPiezo1) or 5 µM Escin (hPIEZO1 and HT1080 WT and P1^KO^) (Sigma-Aldrich, Germany).

For mPIEZO1, the following measures were taken: SC-560: 0.1⍰μM (n⍰=⍰43), 0.3⍰μM (n⍰=⍰41), 1⍰μM (n⍰=⍰40), 3⍰μM (n⍰=⍰40) and 10⍰μM (n⍰=⍰42), OTB: 0.1⍰μM (n⍰=⍰37), 0.3⍰μM (n⍰=⍰34), 1⍰μM (n⍰=⍰29), 3⍰μM (n⍰=⍰29) and 10⍰μM (n⍰=⍰21), BBR: 0.1⍰μM (n⍰=⍰44), 0.3⍰μM (n⍰=⍰36), 1⍰μM (n⍰=⍰48), 3⍰μM (n⍰=⍰44) and 10⍰μM (n⍰=⍰44).

For hPIEZO1, the following measures were taken: SC-560: 0.1⍰μM (n⍰=⍰70), 0.3⍰μM (n⍰=⍰62), 1⍰μM (n⍰=⍰65), 3⍰μM (n⍰=⍰57) and 10⍰μM (n⍰=⍰65), OTB: 1⍰μM (n⍰=⍰44), 3⍰μM (n⍰=⍰38), 10⍰μM (n⍰=⍰47), 20⍰μM (n⍰=⍰36) and 30⍰μM (n⍰=⍰32), BBR: 0.1⍰μM (n⍰=⍰85), 0.3⍰μM (n⍰=⍰78), 1⍰μM (n⍰=⍰76), 3⍰μM (n⍰=⍰82) and 10⍰μM (n⍰=⍰95).

Recording solutions contained (in mM) 110 KF, 10 KCl, 10 NaCl, 10 EGTA, and 10 HEPES, pH 7.2 adjusted with KOH (Internal KF 110, Cat. # 08 3007), and 140 NaCl, 4 KCl, 2 CaCl_2_, 1 MgCl_2_, 5 D-glucose monohydrate, 10 HEPES, pH 7.4 adjusted with NaOH (External Standard, Cat. # 08 3001) supplemented with 0.1% DMSO to match the DMSO concentration of the compound solutions. SC-560, OTB and BBR stock solutions were prepared in DMSO and further diluted in External Standard solution to final serial concentrations of 0.1, 0.3, 1, 3 and 10 µM (up to 30 µM where indicated), or to a final single concentration of 5 µM (for HT1080 WT and P1^KO^ cells recordings). GdCl _3_, a non-selective mechanosensitive ion channel inhibitor, was dissolved in the External Standard solution to a final concentration of 30 µM. External Standard solution containing 0.1% DMSO was used as a negative control.

Mechanical stimulation (M-Stim) of PIEZO1 channels was performed by locally dispensing 20⍰µL of solution onto the cell at a pipetting flow of 110 µL/s, synchronized with triggered current recordings at a holding potential of −80 mV. The M-Stim protocol is well established in cell lines overexpressing PIEZO1 (Murciano et al., 2023, Parsonage et al., 2023), but here it was adapted to record currents from cells with endogenous PIEZO1 expression. For compound evaluation, cells received five (or six, where indicated) consecutive M-Stim additions, followed by 5 min incubation with the compound, and one subsequent M-Stim + compound addition. Control and all concentrations of the compound were tested in parallel on one chip.

Only cells with seal resistance >300 MΩ and stable recordings were included in the analysis. To distinguish PIEZO1-mediated response from baseline noise and to exclude artefactual responses, a well was classified as a “responder” when the peak current amplitude was > 100 pA, and the area under the curve (AUC) was exceeding 20 pA·s. n represents the number of wells that met these thresholds, pre-set as online quality control filters in the PatchControl 384 software (Nanion Technologies GmbH), enabling automated well selection during recordings. The peak current density in the presence of the inhibitor was normalised to the peak current density in the presence of reference only, and the response to the compound was further normalised to the control. IC_50_ values were extracted from the Hill equation. Data were analyzed using DataControl 384 (Nanion Technologies GmbH).

### Whole-cell patch-clamp and poking

Human embryonic kidney cells (HEK293) were cultured in Dulbecco’s modified Eagle’s medium (L0102, Biowest) supplemented with 10% fetal bovine serum (S181H, Biowest) and 1% Penicillin/Streptavidin (15140-122, Gibco) and maintained at 37 ºC with 5% CO_2_.

Cells were seeded in 6-well plates at 70-80% confluency and transfected using Lipofectamine 3000 (Invitrogen) with 1 μg of human Piezo1-pIRES plasmid (a generous gift from Dr Frederick Sachs, Jacobs School of Medicine and Biomedical Sciences, University at Buffalo, NY, USA).

For whole-cell patch-clamp recordings, cells were seeded onto 35 mm plastic Petri dishes previously coated with 6 μg/cm2 Poly-L-Lysine (P6282, Sigma-Aldrich) for 1 h at 37 ºC. Before each experiment, culture medium was replaced with an external solution containing (in mM) 140 NaCl, 2.5 KCl, 1.2 CaCl_2_, 0.5 MgCl_2_, 5 glucose, and 10 HEPES (310 mOsm/L, pH 7.4). The intracellular (pipette) solution contained (in mM) 140 CsCl, 1 EGTA, 10 HEPES, 4 ATP, and 0.3 GTP (300 mOsm/L, pH 7.3). Whole-cell currents were recorded at a holding potential of −80 mV, acquired at 10 kHz, and low-pass-filtered at 1 kHz with an Axopatch 200B amplifier. Mechanical stimulation was delivered using a heat-polished glass micropipette mounted on a piezoelectric positioner and stepper motor controller (E-665, Physik Instruments) synchronized with the acquisition program (pClamp10, Molecular Devices) to displace the pipette in 0.5 μm increments. The inactivation time constant (τ inactivation) was calculated from single-exponential fits of the inactivation phase during 250 ms stimulation periods. Time-to-peak was measured from the onset of mechanical stimulation to the maximum current amplitude. All experiments were performed at room temperature (∼23 °C), 24–48 h after cell seeding.

### Changes in RBCs membrane potential elicited by PIEZO activation

Blood from healthy volunteers was withdrawn upon written informed consent into heparinized vacuum tubes in accordance with the guidelines of the Helsinki Declaration of 1975, as revised in 2008. This work has been approved by the institutional (CNRS) Ethical committee and by the French Ministry for Research (declaration DC-2019-3842). Blood was washed thrice with unbuffered saline (154 mM NaCl, mM KCl, 2 mM CaCl_2_) by centrifugation for 5 min at 2,500 g, the buffy coat and plasma removed then with a final step of 1 minute at 12,000 g, and the packed cells stored at 4 ºC until use.

### Membrane potential estimation

The MBE method was used for the monitoring of membrane potential evolution (Hatem et al., 2023). Briefly, when RBCs are suspended in a nominally buffer-free solution in the presence of the CCCP protonophore (20 µM), changes in extracellular pH reflect membrane potential changes since protons are kept at equilibrium across the membrane. The membrane potential (V_M_) can, thus, be estimated from the equation:

V_M_=61.51⍰mV·(pH_i_-pH_o_) in mV

Due to the high red cell buffer capacity, the intracellular pH (pHi) remains constant (at about 7.2) throughout an experiment and can be estimated as the pH of the solution after lysis with Triton™ X-100 at the end of the experiment. Regarding the experimental procedure, 2900 µL of the experimental solution containing 20 µM of CCCP was heated at 37 °C under constant magnetic stirring. For each experiment, 100 µL of packed RBCs (99% hct) were added, to reach a final cytocrit of 3.3%. All inorganic compounds were added at stock solution 1000X in DMSO. The final DMSO concentration never exceeded 0.3%, a concentration that has no effect on either fluxes or membrane potential. Extracellular pH was measured using a HI1083B pH electrode (HANNA Instruments, France) and a PHM210 pHmeter (Radiometer). Sampling and acquisition were done with an electrode amplifier (EA-BTA, Vernier, USA) at a rate of 1 Hz connected to an AD LABQUEST Mini interface (Vernier, USA) with a resolution of 0.01 pH unit. The data were visualized with Logger Lite software (Vernier, France) and analyzed with Prism v10. At the end of each experiment, Triton X-100™ detergent (1% in 3M NaCl) was added, causing total cell lysis and a resulting solution that attains the intracellular pH.

### Evaluation of compound efficacy on RBCs PIEZO1 activity

To evaluate the efficacy of the different compounds, cells, once injected into the unbuffered solution, were allowed to reach membrane potential equilibrium resulting from proton equilibration mediated by CCCP (typically within less than one minute). The compound to be tested was then injected (110 s after the start of the experiment), followed by stimulation with Yoda1 (312 nM, 210 s). The concentration of 312 nM Yoda1 is sufficient to elicit a calcium influx above the maximal activation threshold of the Gárdos channel (Hatem et al., 2023). The efficacy of the tested compound was assessed by measuring the reduction in the extent of hyperpolarization relative to stimulation with vehicle alone (DMSO) and was expressed as a percentage of inhibition.

### RBC deformability

The effects of OTB, BBR or SC-560 on RBC deformability were assessed after activation of PIEZO1 with Yoda1 (5 μm). RBCs were first incubated with Yoda1 for 5 min at room temperature to induce dehydration, followed by the addition of OTB, BBR or SC-560 for an additional 25 min incubation at room temperature. RBCs were incubated in HEPES-buffered Krebs-Ringer solution (Thermo Scientific, Waltham, MA, USA) at a final hematocrit of 20%. Stock solutions of OTB, BBR, and SC-560 were prepared in dimethyl sulfoxide (DMSO, Sigma Aldrich, Saint Quentin Fallavier, France; 99.9%). A control condition was performed with the vehicle (1% DMSO) and 4 concentrations of each inhibitor were tested (10, 50, 100 and 150 mM). The vehicle volume was matched to the volume of drug solutions to ensure a consistent final DMSO concentration across all conditions. Ektacytometry was used to measure RBC deformability (Elongation Index) at 3 Pa, and at 37 °C, as previously described (Baskurt et al., 2009). The effect of Yoda1 alone was measured at several shear stresses ranging from 03 to 30 Pa. RBCs from healthy human samples were obtained after obtention of the informed consent (Ethics Committee reference number: 2023-A02208-37).

### RBC cation content and volume measurements

Fresh venous blood was obtained by venipuncture from informed patients and healthy volunteers in EDTA collecting tubes. The procedure was done according to the latest declaration of Helsinki (2013) and registered by the Ethic committee CPP Île de France (GR-Ex / N° DC-2016-2618). Experiments were performed 24h after withdrawn. Blood was washed 4 times (800 g, 5 min, 4 °C, swinging rotor) at room temperature in Ringer containing in mM: NaCl 147, KCl 5, MgSO_4_ 1, HEPES 10, buffered with NaOH pH7.4 (320 mOsm). For experiments with Yoda1 to activate PIEZO1 in RBCs, washed RBC suspension was set to ≈25% hematocrit and 0.1 µM ouabain (Sigma-Aldrich), 1 mM CaCl_2_ and 5 mM RbCl were added from concentrated stock solutions. At time zero, 2 µM of Yoda1 (Sigma-Aldrich) was added to cell suspension previously dispatched in Eppendorf tubes containing either 10 µM SC-560 or OTB or BBR. Samples were collected 5 and 15 min after Yoda1 addition and centrifuged for 10 min at 20,000 g and 4 °C. RBC pellet was weighted wet and dry after heating overnight at 80 °C as previously described (Allegrini et al., 2022). Dried pellets were diluted in 5 mL milliQ water overnight 4 °C to extract ions. Na^+^, K^+^ and Rb contents were quantified by flame spectroscopy with a Solaar AA spectrometer (Thermo Fisher Scientific).

### RBC Ca ^2+^measurements

4 µl of RBC washed in Ringer without Ca^2+^ were loaded with 5 µM Fluo4 AM in 500 µl Ringer without Ca^2+^ at 37 °C for 30 min. 25 µl of Fluo4 loaded RBC suspension in 975 µl complete Ringer medium were used to quantify relative changes in free intracellular Ca^2+^ concentrations with a FACS Fortessa BD (BD Bioscience). 10,000 events were analysed and the whole population of RBC was considered. Internal RBC fluorescence was assessed on RBC treated without Fluo4 AM. Data are expressed as a ratio of mean internal fluorescence after the addition of drug (IF) versus mean internal fluorescence at time zero before the addition of drug (IF_0_).

## Results

### Identification of Otenabant as a PIEZO1 inhibitor

Despite a strong rationale for developing PIEZO1 inhibitors, availability of validated selective inhibitors is limited. To identify novel PIEZO1 inhibitors, we established a high-throughput screening assay to quantify PIEZO1-dependent Ca^2+^ entry using the Cal520 calcium probe on the Hamamatsu FDSS µCell platform. The assay involved recording a baseline signal, followed by addition of test compounds for 10 min prior to PIEZO1 activation with the specific agonist Yoda1 in the human promyelocytic cell line PLB-985 (**Fig. 1A**). Yoda1 failed to evoke Ca^2+^ elevations in PIEZO1-deficient cells (P1^KO^) generated by CRISPR/Cas9, validating the specificity of the assay (**Fig. 1A and S1A**). A library (APEXBIO) containing a total of 1,971 compounds was screened in duplicate at 10 µM in a 384-well format. Compounds achieving ≥80% inhibition of the Ca^2+^ response were classified as hits. Notably, the previously reported PIEZO1 inhibitor Benzbromarone was identified as hit (**Fig. 1A**), validating the assay. Twenty hit compounds were then purchased and their ability to prevent Ca^2+^ responses evoked by Yoda2, another PIEZO1 agonist with similar efficacy (**Fig. S1B**) was evaluated across a concentration range to determine IC_50_ values. Among confirmed hits, Benzbromarone (BBR) and Otenabant (OTB) exhibited robust, concentration-dependent inhibition of PIEZO1 activity (**Fig. 1B, C**). For OTB, the initial decrease in fluorescence reflects compound addition, which reduces the signal through dilution. In contrast, BBR induces a dose-dependent increase in fluorescence, consistent with non-specific Ca^2+^ mobilization at high concentrations. BBR has been widely used for the treatment of gout as an uricosuric agent (Heel et al., 1977). OTB (aka CP-945,598) was developed as a selective antagonist of the cannabinoid CB_1_ receptor to treat obesity and metabolic disorders but was discontinued due to severe psychiatric effects of related class compounds.

**Figure 1.**
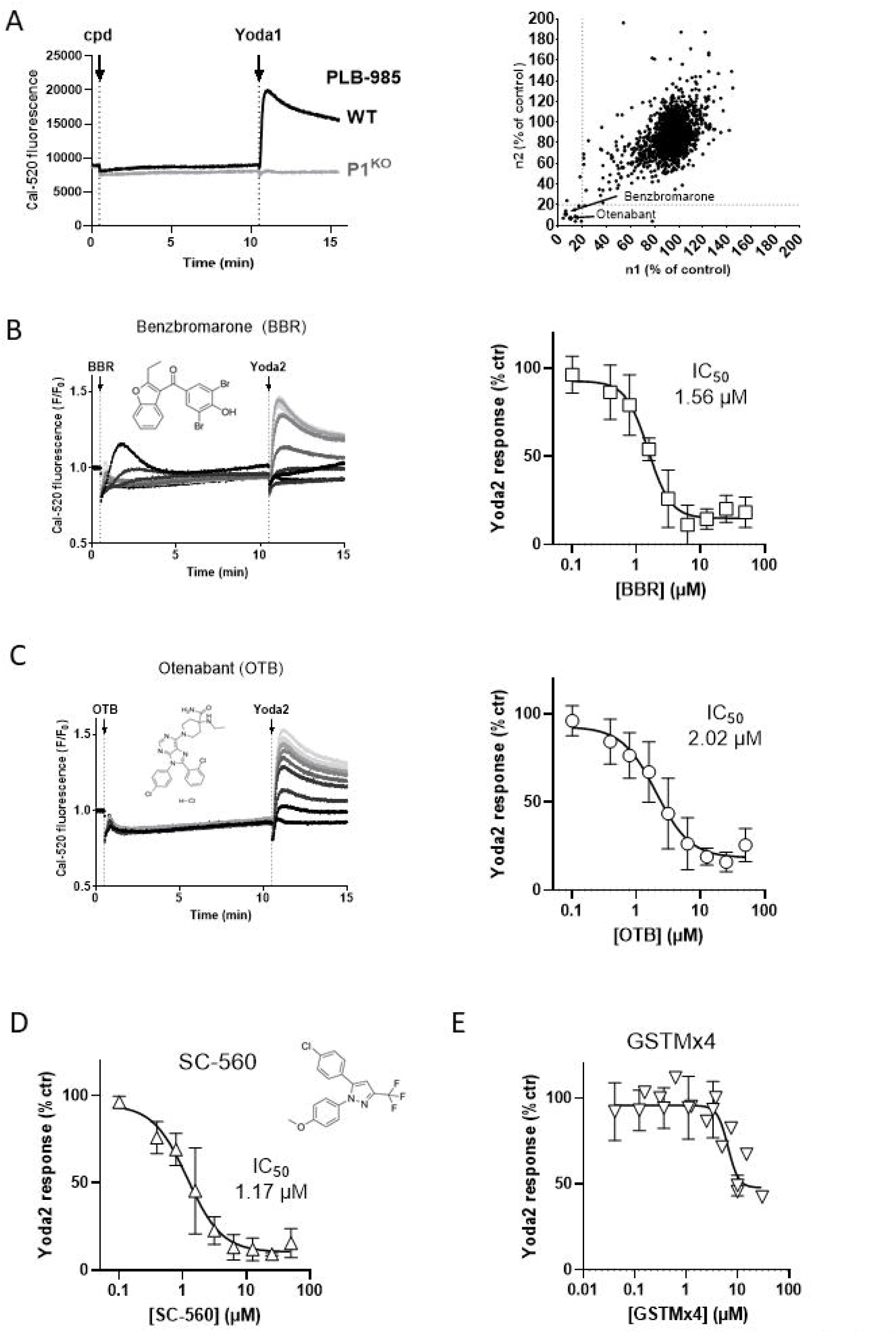
Identification and validation of Otenabant as a PIEZO1 inhibitor. **(A)** Screening assay. Ca^2+^elevations evoked by Yoda1 (5 µM) in wild-type (WT) and PIEZO1-deficient (P1^KO^) human promyelocytic PLB-985 cells exposed to test compounds (cpd) for 10 min were recorded with Cal520 in a Hamamatsu FDSS µCell platform (left). The amplitude of Yoda1-induced Ca^2+^ responses recorded with ∼2’000 tested compounds in two biological replicates was compared (right). Responses were normalized to the DMSO control. Arrows highlight Otenabant (OTB) and Benzbromarone (BBR) as potent inhibitors of the Yoda1-induced Ca^2+^ elevation (>80% inhibition). **(B)** Representative recordings (left) and concentration–response curves (right) of Yoda2-induced Ca_2_ ^+^ elevations at increasing *BBR* concentrations. The chemical structure of BBR is indicated. Increasing compound concentrations are shown from light (0 µM) to dark (50 µM) grey. Note that at high concentrations *BBR* application elicited a transient Ca^2+^ increase. **(C)** Representative recordings (left) and concentration–response curves (right) of Yoda1-induced Ca_2_ ^+^ elevations at increasing OTB concentrations. Chemical structure of OTB is indicated. **(D)** Concentration– response curves of Yoda1-induced Ca^2+^ elevations at increasing SC-560 and **(E)** the tarantula toxin GSTMx4 concentrations. Chemical structure of SC-560 is indicated. Data are mean ±SD of N=5 biological replicates, each measured in technical triplicate.

We next compared the PIEZO1 inhibitory activity of BBR and OTB to 6 reported PIEZO1 inhibitors: the small molecule patented as a PIEZO1 inhibitor SC-560 (JP2014084283 (A)), the tarantula toxin GSMTx4 (Bae et al., 2011), the Yoda1 analogue Dooku-1 (Evans et al., 2018), the cationic inhibitor gadolinium (Gd^3+^) (Coste et al., 2010), and the natural compounds Jatrorrhizine (Hong et al., 2023) and Salvianolic acid B (Pan et al., 2022). SC-560 showed potent inhibition of Yoda2-evoked Ca^2+^ responses in PLB-985 cells (**Fig. 1D**) while GMSTx4 showed a concentration dependent inhibition in its solubility range (< 30 µM) (**Fig. 1E**). Apart from the non-specific trivalent ion Gd^3+^, other claimed PIEZO1 inhibitors were inactive in our assay (**Fig. S1C-F**). We then further compared the newly discovered PIEZO1 inhibitor OTB to the reference compounds BBR and SC-560 in all subsequent experiments.

### 3.2. OTB inhibits human PIEZO1 but not mouse Piezo1

To determine whether the inhibitory effect of OTB on PIEZO1 was cell-type dependent, we assessed its concentration-response profile on Yoda2-induced cytosolic Ca^2+^ elevations in mouse and human cell lines with robust PIEZO1 activity. Using the human promyelocytic cell line PLB-985 as reference, we tested the mouse dendritic cell line JAWSII (**Fig. 2A**), the human fibrosarcoma cell line HT1080 (**Fig. 2B**), and mouse embryonic fibroblasts (MEFs, **Fig. 2C**). To validate the specificity of Yoda2 responses in fibroblasts, we generated PIEZO1-deficient HT1080 cells and Piezo1-deficient MEFs by CRISPR/Cas9A (**Fig 2C, D**). BBR and SC-560 consistently inhibited PIEZO1-dependent Ca^2+^ responses across all tested cell lines, with IC_50_ values in the low micromolar range (**Fig. 2A-C**). In contrast, OTB effectively inhibited PIEZO1-mediated activity in the two human cell lines but had minimal inhibitory activity in the two mouse-derived cell lines, even at high concentrations (**Fig. 2A-C**). To confirm this species selectivity, we used HEK293 cells, which did not intrinsically respond to Yoda2 in our assay and transiently overexpressed either human PIEZO1 or mouse Piezo1. BBR and SC-560 inhibited both the human and mouse orthologs, whereas OTB selectively inhibited hPIEZO1 and was inactive against expressed mPiezo1 (**Fig. 2D**). We then re-expressed a construct encompassing human PIEZO1 and GCamP6 (Yaganoglu et al., 2023) in Piezo1-deficient MEFs to reconstitute hPIEZO1 activity under the control of a doxycycline-dependent promoter in the mouse cellular background (**Fig. S2A**). Yoda2 evoked a small Ca^2+^ response in mP1^KO^ cells reconstituted with hPIEZO1 (mP1^KO^+hP1) in the absence of doxycycline, suggesting a slight leakiness of the inducible promoter, and induced a strong response following doxycycline induction (**Fig. S2B**). OTB did not inhibit endogenous Piezo1 activity in the parental mouse line stably expressing the Cas9 enzyme (**Fig. S2C**) but blocked Yoda2-induced responses in mP1^KO^+hP1 cells (**Fig. S2D**), while BBR and SC-560 similarly inhibited Yoda2 responses in all cases. These data indicate that OTB preferentially inhibits human PIEZO1, potentially reflecting structural differences between the human and mouse mechanosensitive channels.

**Figure 2.**
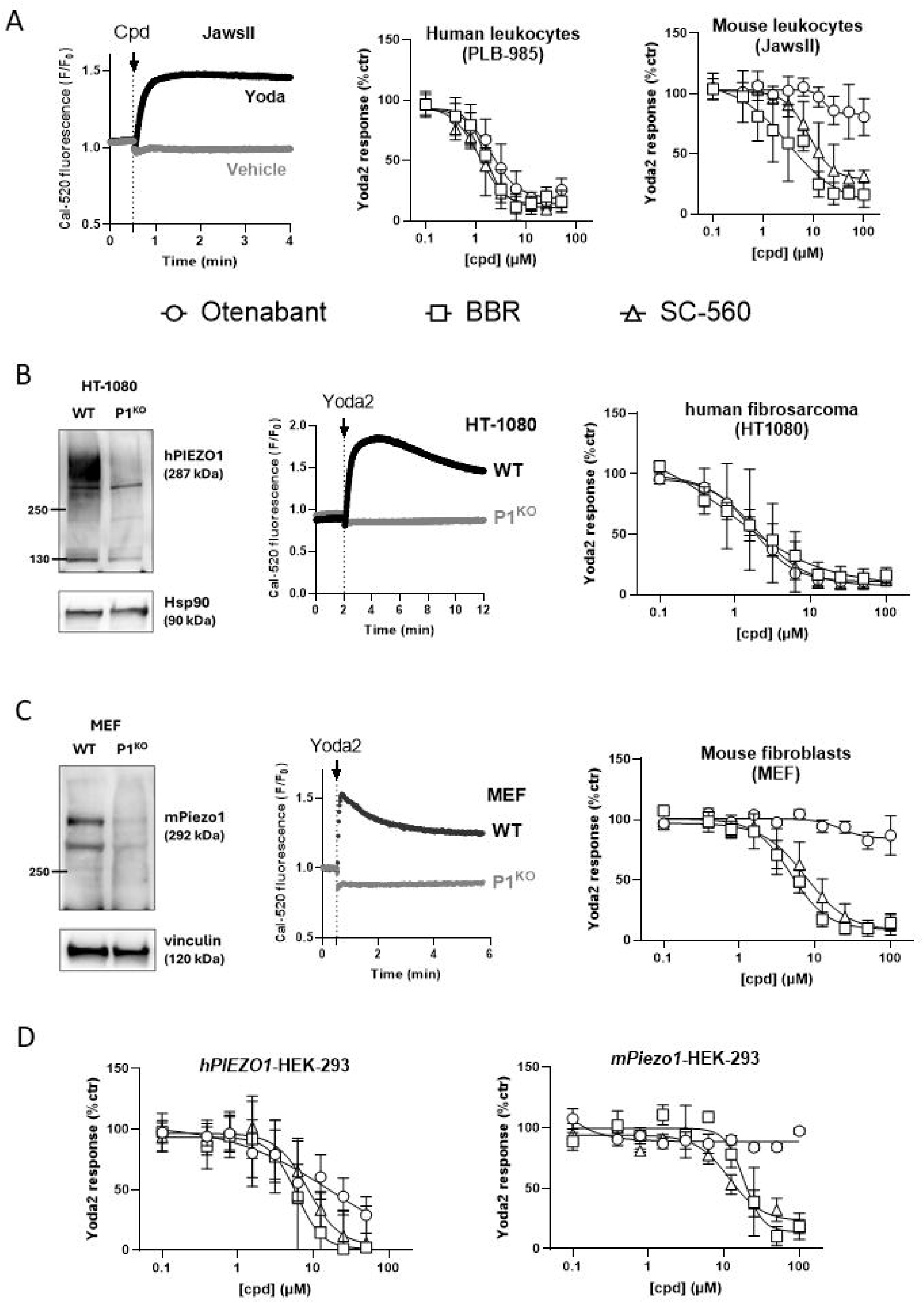
Effect of inhibitors on endogenous and expressed hPIEZO1 and mPiezo1. **(A)** Representative recordings of Ca^2+^ elevation elicited by 5 µM Yoda2 in JawsII cells (left). Concentration–response curves for OTB, BBR and SC-560 on Ca^2+^ elevations evoked by 5 µM Yoda2 in human PLB-985 cells (middle, data from Fig 1 and S1) and mouse JawsII cells (right). OTB showed no inhibitory activity in the mouse dendritic cell line. **(B)** Immunodetection of PIEZO1 in WT and P1^KO^ HT1080 cells by western blot (left). Representative recordings of Ca^2+^ elevation elicited by 5 µM Yoda2 in HT1080 (middle). Concentration–response curves for OTB, BBR and SC-560 on Ca^2+^ elevations evoked by 5 µM Yoda2 in HT1080 (right). OTB showed potent inhibitory activity in the human fibrosarcoma cell line. **(C)** Immunodetection of Piezo1 in WT and P1^KO^ MEF cells by western blot (left). Representative recordings of Ca^2+^ elevation elicited by 5 µM Yoda2 in MEF (middle). Concentration–response curves for OTB, BBR and SC-560 on Ca^2+^ elevations evoked by 5 µM Yoda2 in mouse embryonic fibroblasts (right). OTB showed no inhibitory activity in mouse fibroblasts. **(D)** Concentration–response curves for OTB, BBR and SC-560 in HEK293 cells transiently expressing human PIEZO1 or mouse Piezo1. OTB was inactive against Yoda2-induced activation of the mouse Piezo1 ortholog. Data are mean ±SD of N=3-5 biological replicates, each measured in technical triplicate.

To obtain additional pharmacological insight on the mechanism of PIEZO1 inhibition, we used Jedi2, an alternative PIEZO1 agonist with a mode of action distinct from Yoda2 (Wang et al., 2018). Jedi2 caused a modest Ca^2+^ elevation in PLB-985 cells at high concentrations (1 mM), an effect not observed in PIEZO1-deficient cells, confirming its specificity (**Fig. S2E**). OTB (IC_50_ = 4.7 µM), BBR (IC_50_ = 0.87 µM) and SC-560 (IC_50_ = 3.79 µM) potently inhibited Jedi2-induced PIEZO1 activation, confirming the PIEZO1 inhibitory activity of these compounds (**Fig. S2F**). Interestingly, BBR exhibited the highest potency in this assay, possibly reflecting a different activation mechanism of Jedi2. These data validate OTB as a novel inhibitor of PIEZO1-mediated Ca^2+^ entry evoked by chemical agonists and reveal an unexpected preference for inhibition of the human over the mouse ortholog. To assess a potential effect of OTB on the Yoda2 binding site, we performed a functional competition assay by measuring Ca^2+^ elevations in PLB-985 cells exposed to Yoda2 and OTB mixed at different concentrations. We observed minimal effects on EC_50_ but a dose-dependent decrease in E_max_, suggesting negative allosteric modulation of Yoda2 response (**Fig. S2G**).

### OTB inhibits hPIEZO1 currents activated by shear stress

Mechanically activated currents mediated by mPIEZO1 and hPIEZO1 were recorded using an automated patch-clamp platform (described in (Murciano et al., 2023)) in response to controlled mechanical stimulation (M-Stim).

In mPIEZO1-expressing cells, SC-560 and BBR inhibited mechanically activated inward currents in a concentration-dependent manner (**Fig. 3A, B**). SC-560 produced a gradual reduction in peak current density (IC_50_ = 3.01 µM), while BBR showed higher potency, inhibiting mPIEZO1 currents (IC_50_ = 0.35 µM). In contrast, OTB did not inhibit mPIEZO1-mediated currents, and a reliable IC_50_ value could not be determined within the tested concentration range.

**Figure 3.**
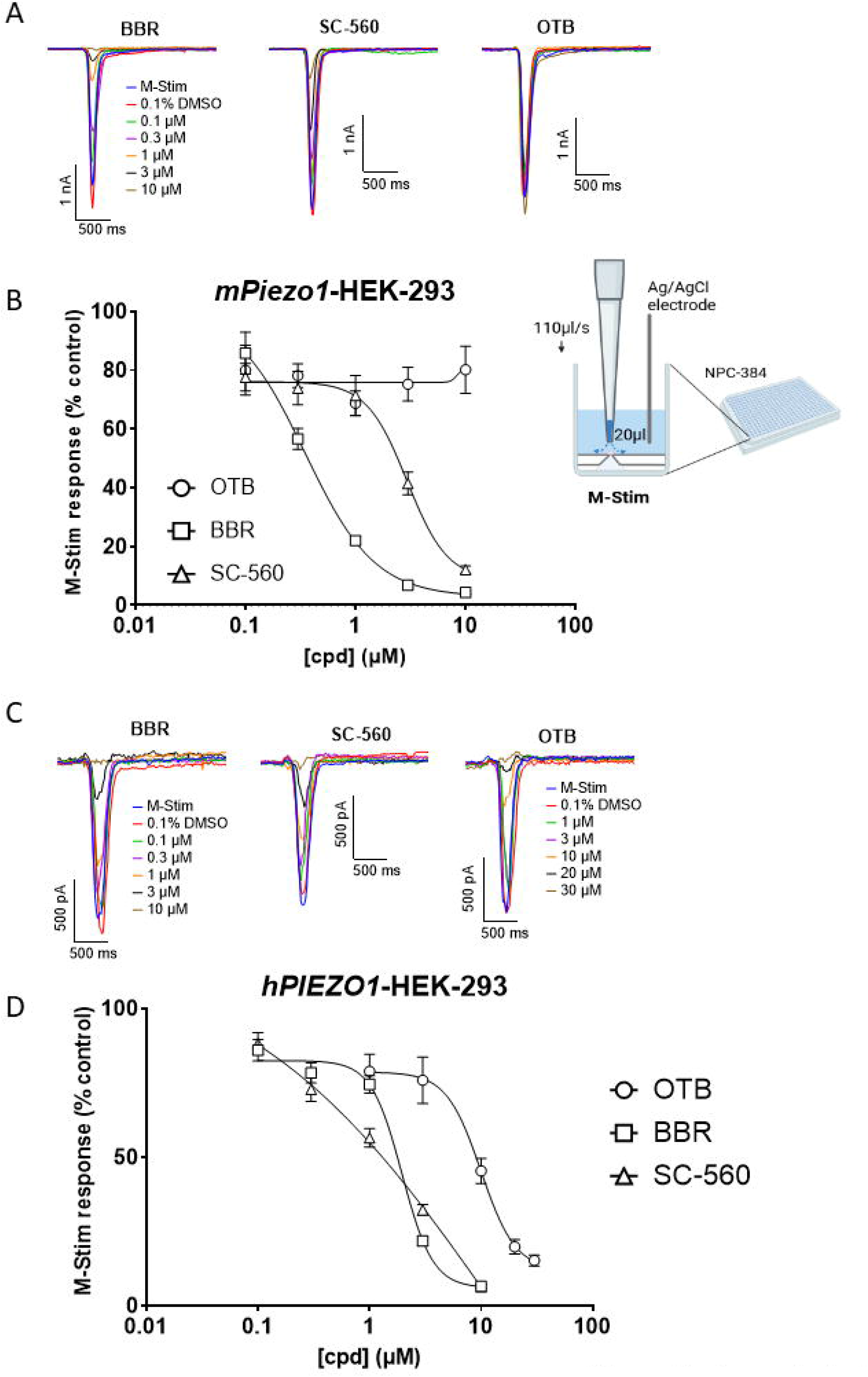
Effect of inhibitors on mechanically activated currents mediated by expressed mPiezo1 and hPIEZO1. **(A)** Representative recordings of mPiezo1-mediated currents evoked by mechanical stimulation (M-Stim) in the absence and presence of BBR, OTB and SC-560. **(B)** Concentration-response curves of M-Stim-evoked mPiezo1 currents recorded at different concentrations of SC-560, OTB and BBR (0.1, 0.3, 1, 3 and 10⍰μM). The sketch shows the mechanical stimulation used in the automated patch-clamp assay. **(C)** Representative recordings of hPIEZO1-mediated currents evoked by M-Stim in the absence and presence of BBR, SC-560 and OTB. **(D)** Concentration-response curves of M-Stim-evoked hPIEZO1 currents recorded at different concentrations of SC-560 and BBR (0.1, 0.3, 1, 3 and 10⍰μM) and OTB (1, 3, 10, 20 and 30⍰μM). The peak current density in the presence of the inhibitor was normalised to the peak current density in the presence of reference only, and the response to the compound was further normalised to the response of 0.1% DMSO used as control. Fitted curves were generated from the Hill equation. Data are mean ±SEM of N=21-95 wells per concentration.

In hPIEZO1-expressing cells, all three compounds inhibited mechanically activated currents in a concentration-dependent manner (**Fig. 3C, D**). SC-560 inhibited hPIEZO1 with potency comparable to mPIEZO1 (IC_50_ = 2.71 µM). OTB also inhibited hPIEZO1-mediated currents, (IC_50_ = 8.78 µM). BBR inhibited hPIEZO1 currents (IC_50_ = 1.96 µM) with greater potency than SC-560 and OTB but reduced potency compared with its effect on mPIEZO1. Across both orthologues, BBR consistently exhibited the greatest inhibitory efficacy, while SC-560 showed species-dependent differences in potency. OTB displayed no inhibition of mPiezo1 and moderate inhibition of hPIEZO1 in this system.

To assess whether OTB inhibits mechanically activated endogenous PIEZO1 currents, we recorded endogenous PIEZO1 activity in the human fibrosarcoma cell line HT1080. M-Stim, applied via repeated high-pressure fluid jets, elicited robust inward currents that were absent in PIEZO1-deficient HT1080 cells and in cells exposed to 30 µM Gd^3+^, confirming that the recorded current was carried by PIEZO1 (**Fig. S3A**). Using this cell model, compounds were tested at a single concentration (5 µM). BBR and OTB potently inhibited the mechanically evoked currents, reducing the total charge carried by 96% and 80%, respectively, whereas SC-560 showed no significant inhibition on endogenous PIEZO1 currents evoked by M-Stim (**Fig. S3B**).

### OTB inhibits hPIEZO1 currents activated by poking

Confirming these data, OTB (30 µM) strongly inhibited inward currents elicited by repeated poking of HEK293 cells transfected with human PIEZO1 (**Fig. 4A, B**). OTB decreased the magnitude of the current in the whole range of poking stimulation and delayed the current inactivation kinetics, suggesting that OTB influences the gating dynamics of PIEZO1 (**Fig. 4C, D**). These data validate OTB as an inhibitor of the PIEZO1 channel independently of its chemical or mechanical mode of activation.

**Figure 4.**
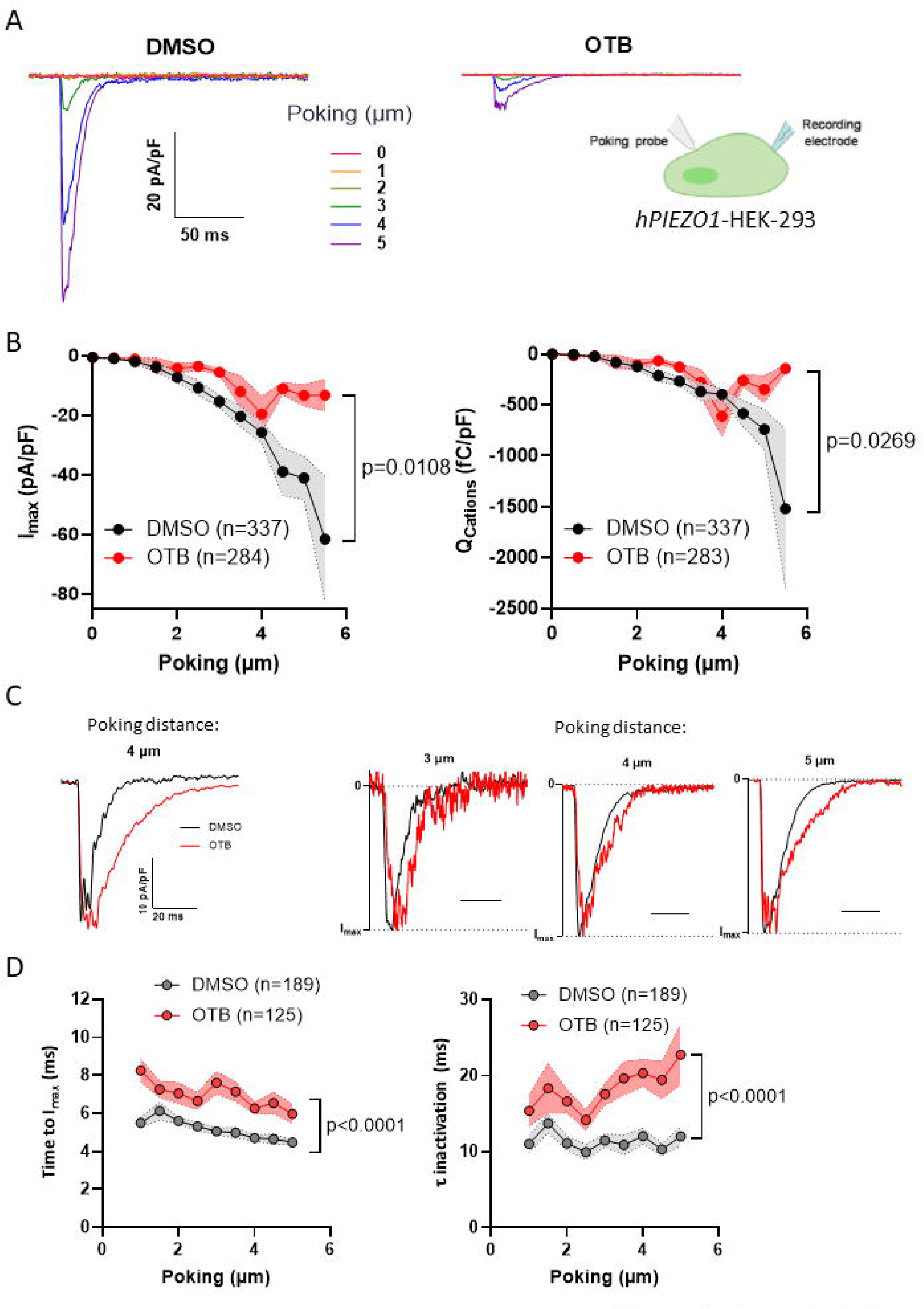
Effect of OTB on hPIEZO1-mediated currents evoked by mechanical poking. **(A)** Representative whole-cell currents recorded at a holding potential of −80 mV in HEK-P1 cells exposed to 0.1 % DMSO (left) or 30 µM OTB (right) during a series of mechanical stimulation with a poking probe (inset), increasing from 0 to 5 µm in 0.5 μm increments. Only traces from 1 μm displacement steps are shown for clarity. **(B)** Peak currents densities (I_max_, left) and cation influx densities (Q_Cations_, right) recorded at increasing displacement steps in HEK-P1 cells treated with DMSO or OTB. N = 46 and 43 cells, One-tailed ratio paired Student’s t test (I_max_) and Wilcoxon matched-pairs signed rank test (Q_Cations_) over the whole range of stimulation steps (N = 11). The total number of episodes recorded is indicated. **(C)** Representative recordings of hPIEZO1-mediated currents elicited by a 4 μm mechanical step in DMSO (black) and OTB (red) illustrating the activation and inactivation kinetics of currents of similar amplitudes (left) and representative recordings of hPIEZO1-mediated currents activated by 3, 4 and 5 µm poking steps, normalized to peak current amplitude (right). N = 46 and 43 cells, One-tailed ratio paired Student’s t test over the 1-5 µm range of stimulation steps (N = 10). The total number of episodes recorded is indicated.

### BBR but not OTB and SC-560 has off-targets effects

Cross-inhibition is common among Ca^2+^ channel inhibitors, and selectivity is often dose-dependent. We therefore tested the inhibitory effect of our compounds on Ca^2+^-permeable channels unrelated to PIEZO1. HL-60 cells, the parental cell line from which PLB-985 cells are derived, express functional purinergic receptors activated by ATP, including ionotropic P2X1 (Buell et al., 1996)and G-protein-coupled receptors P2Y (Communi et al., 2000). ATP (1 µM) induced Ca^2+^ elevations of identical amplitude in WT and P1^KO^ PLB-985 cells (**Fig. 5A**). The ATP-induced Ca^2+^ elevations were not affected by OTB and SC-560 but were efficiently inhibited by *BBR* (IC_50_ = 4.7 µM), indicating a pleiotropic effect of this compound on P2 purinergic receptors (**Fig. 5A**).

**Figure 5.**
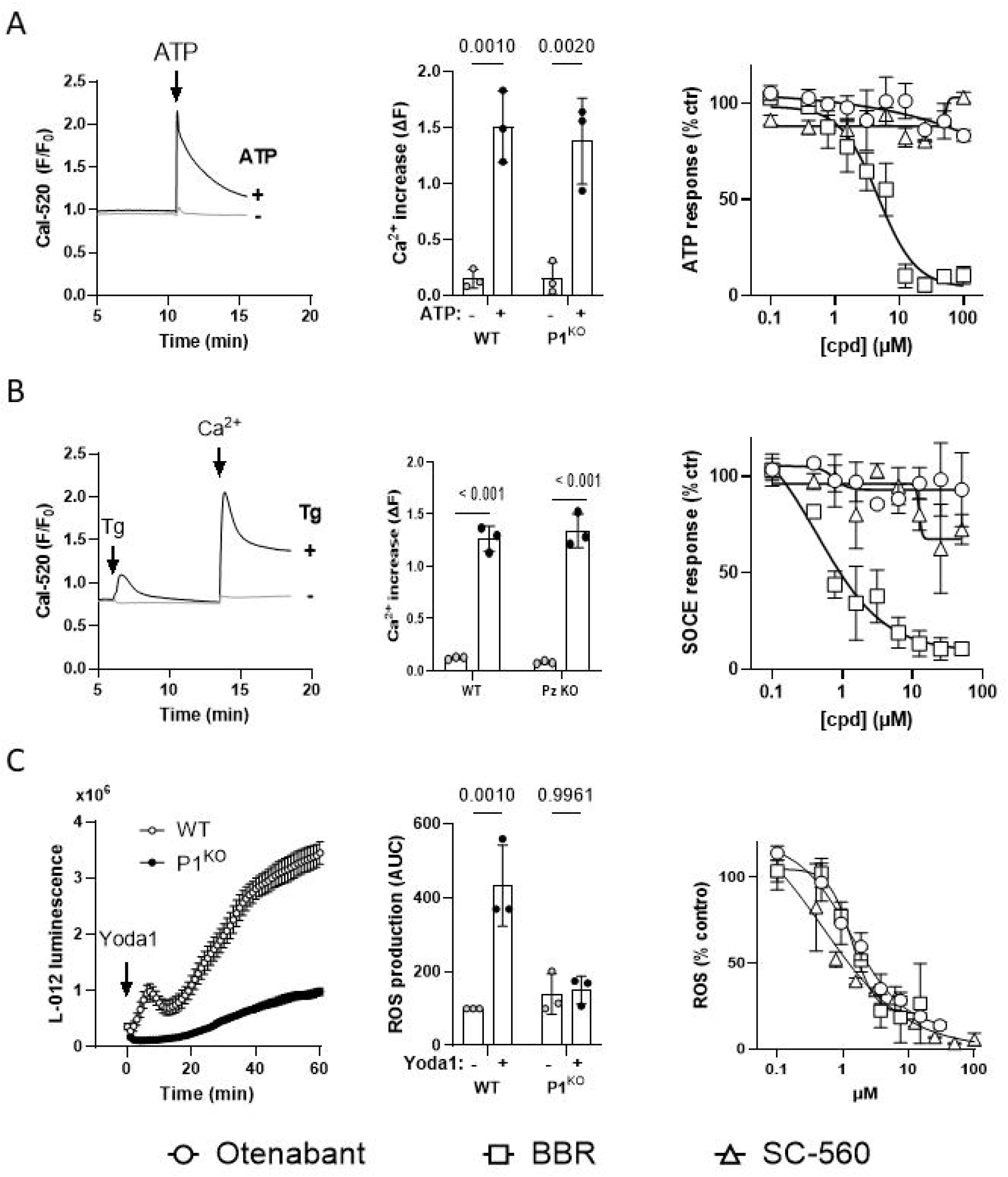
Off-targets effects of PIEZO1 inhibitors and orthogonal assays. **(A)** Representative Ca^2+^ elevation evoked by ATP in PLB-985 cells (left), quantification of ATP-induced Ca^2+^ responses in WT and P1^KO^ cells (middle) and concentration–response curves for OTB, BBR, and SC-560 on ATP-induced Ca^2+^ responses (right). BBR but not OTB or SC-560 dose-dependently inhibited ATP-dependent Ca^2+^ elevations. **(B)** Representative Ca^2+^ elevations evoked by Ca^2+^ readmission to PLB-985 cells treated or not with thapsigargin (Tg) in Ca^2+^-free medium (left), quantification of store-operated Ca^2+^ entry (SOCE) evoked by Tg in WT and P1^KO^ cells (middle) and concentration–response curves for OTB, BBR, and SC-560 on Tg-induced SOCE responses (right). BBR but not OTB or SC-560 dose-dependently inhibited SOCE. **(C)** ROS elevations evoked by Yoda1 in differentiated WT and P1^KO^ PLB-985 cells reported by the chemiluminescent probe L-012 (left), quantification of Yoda1-induced ROS elevations in WT and P1^KO^ cells (middle) and concentration–response curves for OTB, BBR, and SC-560 on Yoda1-induced ROS responses. N=3, two-Way ANOVA, Multiple Comparison with Tukey test.

We next assessed whether the compounds affected the ability of Thapsigargin (Tg) to mobilize Ca^2+^ from intracellular stores and to induce store-operated Ca^2+^ entry (SOCE) mediated by interaction between STIM proteins and Ca^2+^-permeable Orai channels (Yeromin et al., 2006). Addition of Tg in Ca^2+^-free medium and subsequent re-addition of Ca^2+^ evoked Ca^2+^ fluxes of identical amplitude in WT and P1^KO^ PLB-985 cells (**Fig. 5B, middle panel**). The Tg-evoked Ca^2+^ release from intracellular stores and the subsequent Ca^2+^ entry was not impacted by OTB or SC-560 but were both dose-dependently inhibited by BBR (**Fig. 5B and S5A**). Altogether, these results indicate that BBR acts as a promiscuous modulator of Ca^2+^ signalling, exhibiting multiple modes of action on Ca^2+^ channels.

### OTB inhibits PIEZO1-mediated ROS and TNF production

Once differentiated into neutrophil-like cells, PLB-985 cells can generate large amounts of reactive oxygen species (ROS) via the Ca^2+^- and phosphorylation-dependent activation of the phagocytic NADPH oxidase (Zhen et al., 1993). To confirm the activity of the compounds in orthogonal assays, we recorded PIEZO1-dependent oxidant generation with the chemiluminescent probe L-012. Yoda1 induced a detectable production of ROS in differentiated WT but not P1^KO^ PLB-985 cells. ROS generation in PLB-985 cells was potently inhibited by OTB (IC_50_= 1.35 µM), BBR (IC_50_= 1.35 µM), and SC-560 (IC_50_= 0.50 µM), confirming that the inhibitory activity of these compounds persists in orthogonal assays (**Fig. 5C**). We next measured the secretion of TNF by differentiated PLB-985 cells exposed to LPS for 4h and co-stimulated with Yoda1 (10 µM) for 1 h. The Yoda1-induced TNF secretion was absent in P1^KO^ PLB-985 cells and was significantly inhibited by OTB (10 µM) and BBR (10 µM) in WT cells (**Fig. S5B**). These findings establish that PIEZO1 regulate several neutrophil effector functions and confirm that *OTB* and the other PIEZO1 inhibitors can inhibit PIEZO1-regulated cellular processes. In addition, because OTB is a potent CB1R antagonist, we investigated whether CB1R blockade influences PIEZO1 activity by assessing the potential inhibitory effects of several established CB1R inhibitors. In this context, Taranabant, AM251, and AM4113 were completely inactive in Yoda2-treated PLB-985 cells (**Fig. S5C**).

### OTB inhibits PIEZO1-mediated Ca^2+^ influx, hyperpolarization and deformability of red blood cells

PIEZO1 channels alter RBC volume by causing an influx of Ca^2+^ during mechanical stretch that activates the Gárdos Ca^2+^-activated potassium channels (Cahalan et al., 2015). This mechanism plays a role in RBC rheology, as the loss of K^+^ and water enables RBC to deform as they transit through narrow capillaries (Kaestner et al., 2025). Several gain-of-function mutations in *PIEZO1* have been identified in patients suffering from the RBC dehydrating disease hereditary xerocytosis (Jankovsky et al., 2021). To test whether our inhibitors impact this pathway, we measured the changes in membrane potential induced by Yoda1 in RBC from healthy human donors. As previously reported, addition of Yoda1 (312 nM) caused a transient hyperpolarization of RBCs, toward Nernst equilibrium for K^+^ (**Fig. 6A, left**). This hyperpolarization was prevented by pre-incubation with the K^+^ channel blocker charybdotoxin (100⍰nM), confirming that it was solely elicited by the increased activity of Gárdos channels whose opening probability strictly depends on the intracellular Ca^2+^ concentration (**Fig. S6A**). Preincubation of RBC for 100 s with increasing concentrations of OTB, BBR, and SC-560 dose-dependently inhibited the Yoda1-induced RBC hyperpolarization (**Fig. 6A, right**). These results indicate that OTB and the other PIEZO1 inhibitors inhibit the hyperpolarization of red blood cells caused by chemical activation of PIEZO1 and the resulting Ca^2+^ activation of Gárdos channels.

**Figure 6.**
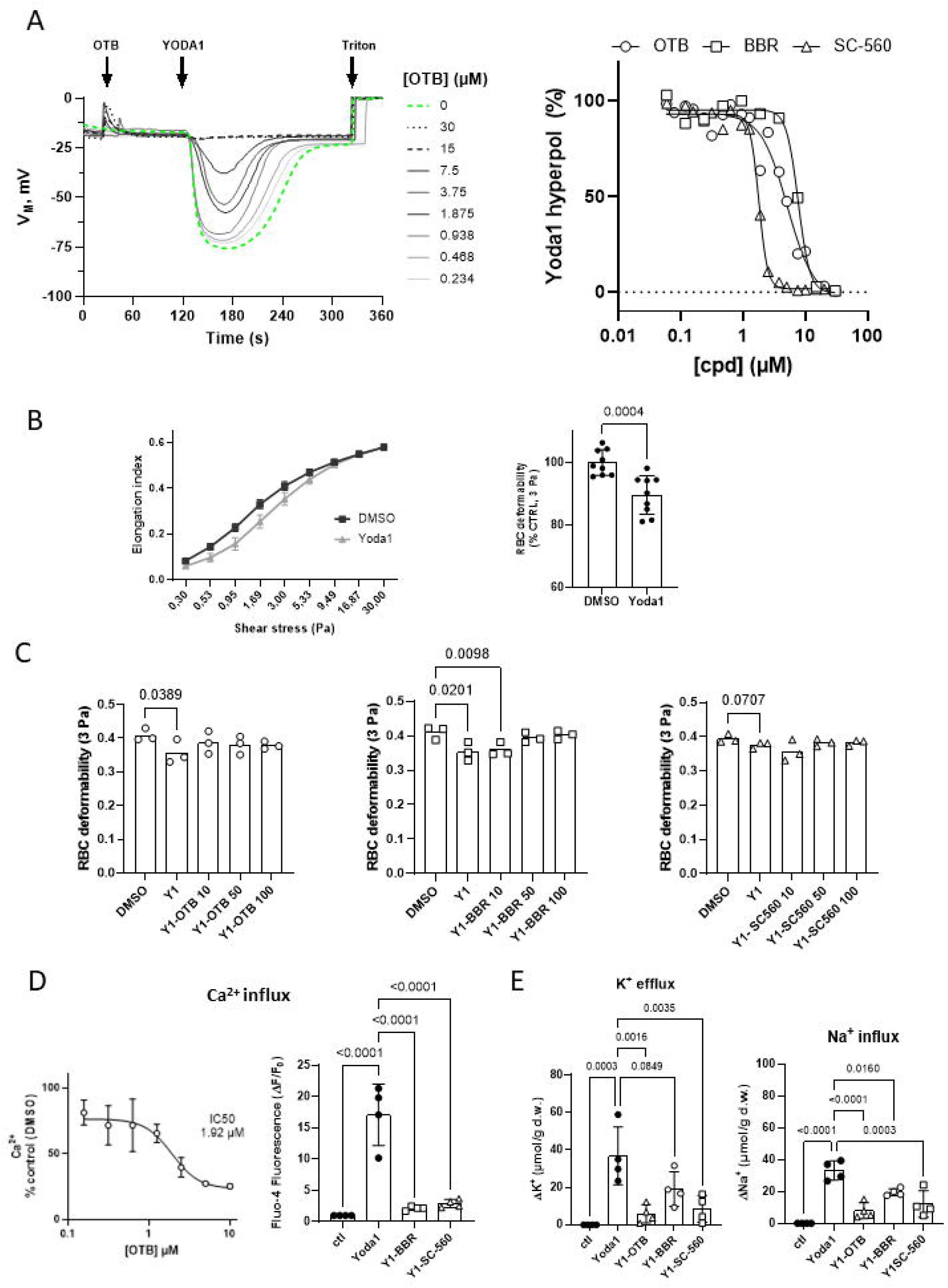
Otenabant inhibits PIEZO1-mediated ion fluxes and volume changes in human red blood cells. **(A)** Representative recordings of the membrane potential changes induced by Yoda1 (312 nM) in human red blood cells (RBCs) exposed to increasing concentration of OTB. Dashed green line indicates the control response in DMSO (left). Concentration–response curves for OTB, BBR and SC-560 on Yoda1-induced depolarization of RBCs. N = 3, 2, and 1 recording for OTB, SC-560, and BBR, respectively (right). (**B**) Representative elongation curve at different shear stress of RBCs treated with DMSO or 5 µM Yoda1 (left). Quantification of RBC deformability at 3 Pa in DMSO or Yoda1 (right). Data are mean ± SD, one-way ANOVA with Tukey’s multiple comparisons test; n = 9. (**C**) Deformability at 3 Pa of RBCs treated with Yoda1 and different concentrations of OTB, BBR and SC-560. **P < 0.01, vs control, Friedman test with Dunn’s post-hoc test; n = 3. (**D**) Concentration–response curves for OTB on Yoda1-induced influx of Ca^2+^ influx (left) and variation of Influx of (right) evoked by Yoda1 (2 µM) in RBC pre-treated or not with 10 µM BBR and SC-560. (**E**) Variation of K^+^ efflux, absolute values (left) and of Na^+^ influx (right) evoked by Yoda1 (2 µM) in RBC pre-treated or not with 10 µM OTB, BBR and SC-560 (control values for K^+^ and Na^+^ were 219.8±8.9 and 14.1±6.9 µmol/g d.w. respectively). Data are mean ± SD, one-way ANOVA with Tukey’s multiple comparisons test; n = 3-4 independent experiments, each measured in triplicate.

We then assessed RBC deformability by ektacytometry, a laser diffraction–based technique routinely applied in the diagnosis of hereditary xerocytosis and other RBC disorders. Treatment of RBCs with Yoda1 resulted in a significant reduction in RBC deformability (**Fig. 6B**), to levels comparable to those observed in RBC from patients with hereditary spherocytosis (Casabianca et al., 2024). Subsequent addition of OTB, BBR, or SC-560 partially rescued the Yoda1-induced loss of deformability (**Fig. 6C**).

We next evaluated the effects of the inhibitors on the ion fluxes evoked by PIEZO1 activation in RBC. To prevent Yoda1-induced dehydration, Senicapoc, a selective blocker of the Gàrdos Ca^2+^-dependent K^+^ channel, was added to the cell suspension. Consistent with previous observation (Rapetti-Mauss et al., 2017), Yoda1 markedly increased cation passage through PIEZO1. A rapid and robust intracellular Ca^2+^ increase was observed within the first min after Yoda1 addition that was dose-dependently decreased by pre-treatment with the OTB (**Fig. 6D, left**): This inhibitory effect was also observed with 10 µM SC-560 and BBR (**Fig. 6D, right**). Yoda1 treatment triggered a substantial K^+^ efflux (37 µmol/g dry weight over 15 min), which was compensated by a comparable Na^+^ influx (33 µmol/g dry weight over 15 min) (**Fig. 6E**). Pre-incubation with OTB, BBR, and SC-560 significantly reduced both PIEZO1-mediated K^+^ efflux and Na^+^ uptake (**Fig. 6E**).

Finally, we assessed the effects of the compounds in RBC from a patient carrying the *PIEZO1* gain-of-function variant causing hereditary xerocytosis. As expected, the R2456H mutation was associated with a higher basal and Yoda1-induced cation passage compared with RBC from control patients (**Fig. S6B-E**). SC-560, OTB, and BBR reduced PIEZO1-mediated ion fluxes, with SC-560 and OTB displaying greater efficacy than BBR in all assays inhibiting K^+^ efflux and Na^+^ uptake by 60-70%, whereas BBR produced a more modest inhibition (**Fig. S6B-E**).

## Discussion and Conclusions

The pivotal role of the mechanoreceptor PIEZO1 in diverse physiological and pathological processes provides a strong rationale for identifying specific, potent, and well-tolerated inhibitors. Several small molecules have been reported to inhibit PIEZO1, but few have undergone comprehensive evaluation, and many suffer from limited specificity. To date, the tarantula toxin GsMTx4 remains the reference PIEZO1 inhibitor; however, this ∼4-kDa amphipathic peptide has limited utility to selectively study PIEZO1 activity as GsMTx4 acts indirectly via membrane lipids (Cox and Gottlieb, 2019), exhibits poor solubility, and is associated with substantial off-target effects.

Using a Yoda1-based high-throughput screening strategy, we identified the small molecule OTB (CP-945,598) as a previously unrecognized and robust inhibitor of human PIEZO1. OTB is a potent and selective cannabinoid CB_1_ receptor antagonist (Griffith et al., 2009) originally developed by Pfizer for obesity. Although it progressed to phase 3 clinical trials and produced significant dose-dependent weight loss, its development was discontinued due to concerns regarding the psychiatric adverse effects observed across the CB_1_ antagonist class rather than molecule-specific toxicity (Aronne et al., 2011).

OTB consistently inhibited PIEZO1 activity across multiple orthogonal assays, including Yoda1- and 2-induced Ca^2+^ responses, whole-cell electrophysiological recordings in response to mechanical stimulation, and functional measurements in differentiated neutrophil-like cells and human RBCs. The convergence of these independent methods strongly supports a direct effect on PIEZO1 rather than indirect modulation. Importantly, indirect contributions from CB_1_ receptor inhibition are unlikely, as structurally distinct and well-validated CB_1_ antagonists failed to inhibit PIEZO1.

One of the most striking findings of this study is that OTB inhibited human but not mouse PIEZO1, revealing an unexpected species-specific pharmacology. This differential sensitivity suggests structural divergence that may help localize the OTB binding site, for example, by integrating comparative cryo-EM modelling or domain-swapping mutational approaches. Notably, recent structural comparisons have highlighted meaningful differences between human and mouse PIEZO1, including a more flattened architecture and wider pore radius in the human channel (Shan et al., 2025). However, this species specificity poses challenges for *in vivo* validation, as standard mouse models will likely not accurately reflect human PIEZO1 pharmacology or the activity of OTB-derived inhibitors. Therefore, further evaluations of OTB as a Piezo1 antagonist for in vivo applications will foster alternative approaches to animal experiments. Systems taking advantage of the easy accessibility – also in large numbers – of RBCs as human model cells (Kaestner and Minetti, 2017) may play a key role in such an evaluation concept. Nowadays, even genetically modified RBCs become available in large numbers (Kaestner et al., 2024) and this may allow for a tuned test system based on primary human cells. We did not pursue an in-depth analysis of OTB’s inhibitory mechanism, but our data excluded a direct competitive interaction with Yoda2. Furthermore, the fact that OTB inhibited a reconstituted human PIEZO1 activity in mouse fibroblasts indicates that OTB specificity for human PIEZO1 is not due to a specific human regulatory mechanism. In whole-cell patch-clamp experiments, OTB reduced PIEZO1 current amplitude yet consistently delayed channel inactivation. This slowed inactivation suggests that OTB stabilizes a non-inactivated conformational state of human PIEZO1 while still limiting overall channel activity. A definitive understanding of OTB’s binding site and mechanism of action will require radiolabelled binding studies together with integrated structural, computational, and mutational analyses.

This study also provides the first direct experimental comparison of previously reported PIEZO1 inhibitors confirming that Gd^3+^, GsMTx4, BBR, and SC-560 all inhibited PIEZO1 in our assays. BBR was introduced more than 50 years ago as a uricosuric agent for hyperuricemia and gout, and was subsequently shown to interact with the Urate Transporter 1 (Iwanaga et al., 2005). BBR was withdrawn from European markets in 2004 due to hepatotoxicity concerns (Jansen et al., 2004). BBR was recently identified as a PIEZO1 inhibitor and proposed for repurposing in hereditary xerocytosis (Liang et al., 2024) and sickle cell disease (Liang et al., 2025). Its appearance as a potent hit in our screen validated our screening strategy, but our results revealed substantial off-target effects across several Ca^2+^ channels, strongly limiting its specificity as a PIEZO1 inhibitor in therapeutic applications.

SC-560 was originally derived from celecoxib and has widely been used as a COX-1–selective inhibitor (Smith et al., 1998). SC-560 also inhibited both human and mouse PIEZO1 in our assays. Interestingly, SC-560 was inactive in shear-stress-activated human whole-cell patch-clamp recordings in HT1080 cells, suggesting that its inhibitory activity may be specific to Yoda1-induced gating in HT1080 cells. Nevertheless, SC-560 provides a valuable pharmacological tool for dissecting Yoda1-dependent activation mechanisms.

The identification of OTB as a selective human PIEZO1 inhibitor expands the availability of pharmacological tools for studying mechanosensitive PIEZO1 channels. Beyond its experimental utility, OTB represents a promising chemical starting point for translational exploration. The efficacy of OTB on inhibiting PIEZO1 in RBC and its ability to reverse the deleterious effect of Yoda1 on RBC deformability could be of interest in the context of diseases where RBC deformability is severely reduced, such as hereditary spherocytosis (Casabianca et al., 2024) or sickle cell disease (Tripette et al., 2009). The sub-micromolar potency of OTB and prior clinical evaluation provide a strong foundation for medicinal chemistry programs aimed at improving specificity while avoiding CB1 receptor activity. Future optimization should prioritize decoupling the molecule from the CB1 receptor pharmacophore space to advance OTB-derived compounds as genuine PIEZO1 inhibitors.

In summary, our findings demonstrate that Yoda-based functional screening is an effective strategy for identifying small-molecule PIEZO1 antagonists and revealed OTB as a potent, species-selective inhibitor of human PIEZO1. These results broaden the pharmacological landscape of PIEZO1 modulation and provide a foundation for next-generation PIEZO1-targeted therapeutics. Continued work integrating structural studies, medicinal chemistry, and *in vivo* validation will be essential for unlocking the full therapeutic potential of this new inhibitory scaffold.

## Author Contributions

Vincent Jaquet: experiment design, supervision, data acquisition, data analysis, writing; Reetta Penttinen: experiment design, data acquisition, data analysis and visualisation; Gorane Rodríguez-Urquirizar: data acquisition, data analysis, and editing;

Cyril Castebou: data acquisition;

Yves Cambet: data acquisition, data analysis;

Nicolas Rosa: data acquisition, data analysis, and editing;

Mathilde Bourdin: data analysis and editing;

Bita Asghariastanehei: data acquisition, data analysis, and editing;

Franciele de Lima: data acquisition, data analysis, and editing;

Marie Martin: data acquisition, data analysis, and editing;

Elie Nader: data analysis and editing;

Selma A. Serra: experiment design, data acquisition, data analysis, and editing;

José M. Fernández-Fernández: funding acquisition, experiment design, supervision, data analysis, and editing;

Nicoletta Murciano: experiment design, writing-editing, supervision;

Philippe Connes: data analysis and editing;

Stéphane Egée: data analysis and editing;

Hélène Guizouarn: Ca^2+^, Na^+^, K^+^ measurements in human RBC, WT or with PIEZO1 mutation; data analysis;

Lars Kaestner: funding acquisition, supervision, and editing; Niels Fertig: funding acquisition and proof-reading;

Maria Giustina Rotordam: experiment design, supervision, and editing;

Nicolas Demaurex: experiment design, supervision, funding acquisition and editing.

## Acknowledgements

This work was funded by the Swiss National Foundation (grant number 310030_219547 to ND), by the European Community (Marie Skłodowska-Curie project no. 101120168— INNOVATION), by the German Research Foundation (DFG), project number 522062907, by the Spanish Ministry of Science and Innovation (MCIN)/State Research Agency (AEI, Agencia Estatal de Investigación)/10.13039/501100011033/, and FEDER Funds (Fondo Europeo de Desarrollo Regional) (grant PID2022-136546OB-I00 to J.M.F-F); “Unidad de Excelencia María de Maeztu” CEX2024-001431-M, funded by MICIU/AEI/10.13039/501100011033. HG was supported by the France 2030 program through the IDEX Paris Cité InIdex GR-Ex.

HEK T-REx™ 293 cells overexpressing hPIEZO1 and mPiezo1 for automated patch-clamp recordings were kindly provided by Prof. David J Beech (University of Leeds, Leeds, UK).

## FIGURE LEGENDS

<u>Figure S1</u>. **Evaluation of reported PIEZO1 inhibitors**.

**(A)** Immunodetection of PIEZO1 in WT and P1^KO^ PLB-985 cells by western blot. **(B)** Representative recordings of Ca^2+^ elevations elicited by 5 µM Yoda1 and Yoda2 in PLB-985 cells (left) and concentration–response curves (right), indicating an EC_50_ of 2.03 µM for Yoda1 and of 1.3 µM for Yoda2. **(C-F)** Concentration–response curves for *Dooku1* (C), Gadolinium (Gd^3+^) (D), Jatrorrhizine (E), and Salvianolic acid (F) on Ca^2+^ elevations evoked by 5 µM Yoda2 in PLB-985 cells. Data are mean ±SD of N=3 biological replicates, each measured in technical triplicate.

<u>Figure S2</u>. **Effect of inhibitors on hPIEZO1 expressed in Piezo1-deficient MEF and on Jedi2 responses**.

**(A)** Western blot showing PIEZO1 immunoreactivity in Piezo1-deficient MEF cells (mP1^KO^) and mP1^KO^ cells stably expressing human PIEZO1 driven by a doxycycline promoter (*hPIEZO1*-*mP1*^*KO*^). **(B)** Ca^2+^ responses elicited by 5 µM Yoda2 in *hPIEZO1*-*mP1*^*KO*^ before and after addition of doxycycline., revealing a minor leakiness of the inducible promoter. **(C, D)** Concentration–response curves for OTB, BBR and SC-560 in MEF^*Cas9*^ cells and in *hPIEZO1*-*mP1*^*KO*^. *OTB* had no inhibitory activity in mouse cells expressing endogenous mPiezo1 but inhibited Yoda2-induced activity in knock-out cells reconstituted with hPIEZO1. **(E, F)** Ca^2+^ elevations evoked by 1 mM Jedi2 in WT and PIEZO1-deficient PLB-985 cells and concentration–response curves for OTB, BBR and SC-560 on Ca^2+^ elevations evoked by 1 mM Jedi2 in PLB-985 cells. **(G)** Quantification of a potential competition between OTB and Yoda2. The profile of inhibition of Yoda2-induced Ca^2+^ elevation is compatible with a non-competitive inhibition. Data are mean ±SD of N=2-4 biological replicates, each measured in technical triplicate.

<u>Figure S3</u>. **Effect of inhibitors on mechanically activated endogeneous hPIEZO1 currents**.

**(A)** Representative recordings of M-Stim-evoked currents in wild-type (WT) and PIEZO1-edited (P1^KO^) HT1080 cells before and after exposure to 30 µM Gd^3+^ (left, black and red traces, respectively). Peak current densities and total charge displaced by M-Stim in WT cells (right, N=7). **(B)** Representative recordings (left), peak current densities (middle) and total charge displaced (right) by M-Stim in WT cells before and after exposure to the indicated compounds (all at 5 µM in 0.1 % DMSO). N = 8, 8, 9, and 7 wells for DMSO, SC-560, OTB, and BBR, respectively, ratio paired Student’s t test (DMSO, SC-560, OTB) and Wilcoxon test (Gd^3+^, BBR).

<u>Figure S5</u>. **Off-targets effects of PIEZO1 inhibitors, orthogonal assays and mode of action**

**(A)** Representative Ca^2+^ elevation evoked by Tg in PLB-985 cells in Ca^2+^-free medium (left), quantification of the amount of Ca^2+^ released by Tg in WT and P1^KO^ cells (middle) and concentration–response curves for *OTB, BBR*, and *SC-560* on Tg-induced Ca^2+^ release (right). N=3, two-Way ANOVA, Multiple Comparison with Tukey test. BBR but not OTB or SC-560 dose-dependently inhibited Tg-induced Ca^2+^ release. **(B)** Quantification of TNF secretion by differentiated PLB-985 cells following LPS activation. Yoda1-induced TNF secretion was absent in PIEZO1-deficient cells and abolished by pre-treatment with 10 µM of *OTB* and *BBR*. N=3. **(C)** Concentration–response curves of the CB1R antagonists Taranabant, AM251 and AM4113 on Yoda2-induced responses in PLB-985 cells. Data are mean ±SD of N=2 biological replicates, each measured in technical duplicate.

**<u>Figure S6</u>. Otenabant inhibits PIEZO1-mediated ion fluxes in RBCs from a XH patient**

(**A**) Representative recordings of the membrane potential changes induced by Yoda1 (312 nM) in human red blood cells (RBCs) ± charybdotoxin (100nM). Effect of 10 µM OTB, BBR and SC-560 on (**B)** Ca^2+^ influx variations, Rb^+^ influx (**C)**, K^+^ efflux variations, absolute values (**D**), and Na^+^ influx variations (**E**) evoked by Yoda1 (2 µM) in RBC from a HX patient measured in triplicate.

## References

Allegrini, B., Jedele, S., David Nguyen, L., Mignotet, M., Rapetti-Mauss, R., Etchebest, C., Fenneteau, O., Loubat, A., Boutet, A., Thomas, C., Durin, J., Petit, A., Badens, C., Garcon, L., Da Costa, L. & Guizouarn, H. 2022. New KCNN4 Variants Associated With Anemia: Stomatocytosis Without Erythrocyte Dehydration. Front Physiol, 13, 918620.

Aronne, L. J., Finer, N., Hollander, P. A., England, R. D., Klioze, S. S., Chew, R. D., Fountaine, R. J., Powell, C. M. & Obourn, J. D. 2011. Efficacy and safety of CP-945,598, a selective cannabinoid CB1 receptor antagonist, on weight loss and maintenance. Obesity (Silver Spring), 19, 1404–14.

Bae, C., Sachs, F. & Gottlieb, P. A. 2011. The mechanosensitive ion channel Piezo1 is inhibited by the peptide GsMTx4. Biochemistry, 50, 6295–300.

Baskurt, O. K., Boynard, M., Cokelet, G. C., Connes, P., Cooke, B. M., Forconi, S., Liao, F., Hardeman, M. R., Jung, F., Meiselman, H. J., Nash, G., Nemeth, N., Neu, B., Sandhagen, B., Shin, S., Thurston, G., Wautier, J. L. & INTERNATIONAL EXPERT PANEL FOR STANDARDIZATION OF HEMORHEOLOGICAL, M. 2009. New guidelines for hemorheological laboratory techniques. Clin Hemorheol Microcirc, 42, 75–97.

Blythe, N. M., Muraki, K., Ludlow, M. J., Stylianidis, V., Gilbert, H. T. J., Evans, E. L., Cuthbertson, K., Foster, R., Swift, J., Li, J., Drinkhill, M. J., Van Nieuwenhoven, F. A., Porter, K. E., Beech, D. J. & Turner, N. A. 2019. Mechanically activated Piezo1 channels of cardiac fibroblasts stimulate p38 mitogen-activated protein kinase activity and interleukin-6 secretion. J Biol Chem, 294, 17395–17408.

Buell, G., Michel, A. D., Lewis, C., Collo, G., Humphrey, P. P. & Surprenant, A. 1996. P2X1 receptor activation in HL60 cells. Blood, 87, 2659–64.

Cahalan, S. M., Lukacs, V., Ranade, S. S., Chien, S., Bandell, M. & Patapoutian, A. 2015. Piezo1 links mechanical forces to red blood cell volume. Elife, 4.

Carreras-Sureda, A., Zhang, X., Laubry, L., Brunetti, J., Koenig, S., Wang, X., Castelbou, C., Hetz, C., Liu, Y., Frieden, M. & Demaurex, N. 2023. The ER stress sensor IRE1 interacts with STIM1 to promote store-operated calcium entry, T cell activation, and muscular differentiation. Cell Rep, 42, 113540.

Casabianca, M., Gauthier, A., Nader, E., Cannas, G., Martin, F., Martin, M., Carin, R., Boisson, C., Guillot, N., Merazga, S., Renoux, C., Bertrand, Y., Garnier, N., Hot, A., Muniansi, I., Halfon-Domenech, C., Poutrel, S., Joly, P. & Connes, P. 2024. Red blood cell senescence and vascular function in patients with hereditary spherocytosis with and without splenectomy. Br J Haematol, 204, e41–e44.

Communi, D., Janssens, R., Robaye, B., Zeelis, N. & Boeynaems, J. M. 2000. Rapid up-regulation of P2Y messengers during granulocytic differentiation of HL-60 cells. FEBS Lett, 475, 39–42.

Coste, B., Mathur, J., Schmidt, M., Earley, T. J., Ranade, S., Petrus, M. J., Dubin, A. E. & Patapoutian, A. 2010. Piezo1 and Piezo2 are essential components of distinct mechanically activated cation channels. Science, 330, 55–60.

Cox, C. D. & Gottlieb, P. A. 2019. Amphipathic molecules modulate PIEZO1 activity. Biochem Soc Trans, 47, 1833–1842.

Evans, E. L., Cuthbertson, K., Endesh, N., Rode, B., Blythe, N. M., Hyman, A. J., Hall, S. J., Gaunt, H. J., Ludlow, M. J., Foster, R. & Beech, D. J. 2018. Yoda1 analogue (Dooku1) which antagonizes Yoda1-evoked activation of Piezo1 and aortic relaxation. Br J Pharmacol, 175, 1744–1759.

Giry-Laterriere, M., Cherpin, O., Kim, Y. S., Jensen, J. & Salmon, P. 2011. Polyswitch lentivectors: “all-in-one” lentiviral vectors for drug-inducible gene expression, live selection, and recombination cloning. Hum Gene Ther, 22, 1255–67.

Gnanasambandam, R., Ghatak, C., Yasmann, A., Nishizawa, K., Sachs, F., Ladokhin, A. S., Sukharev, S. I. & Suchyna, T. M. 2017. GsMTx4: Mechanism of Inhibiting Mechanosensitive Ion Channels. Biophys J, 112, 31–45.

Griffith, D. A., Hadcock, J. R., Black, S. C., Iredale, P. A., Carpino, P. A., Dasilva-Jardine, P., Day, R., Dibrino, J., Dow, R. L., Landis, M. S., O’Connor, R. E. & Scott, D. O. 2009. Discovery of 1-[9-(4-chlorophenyl)-8-(2-chlorophenyl)-9H-purin-6-yl]-4-ethylaminopiperidine-4-carboxylic acid amide hydrochloride (CP-945,598), a novel, potent, and selective cannabinoid type 1 receptor antagonist. J Med Chem, 52, 234–7.

Hatem, A., Poussereau, G., Gachenot, M., Peres, L., Bouyer, G. & Egee, S. 2023. Dual action of Dooku1 on PIEZO1 channel in human red blood cells. Front Physiol, 14, 1222983.

Heel, R. C., Brogden, R. N., Speight, T. M. & Avery, G. S. 1977. Benzbromarone: a review of its pharmacological properties and therapeutic use in gout and hyperuricaemia. Drugs, 14, 349–66.

Hong, T., Pan, X., Xu, H., Zheng, Z., Wen, L., Li, J. & Xia, M. 2023. Jatrorrhizine inhibits Piezo1 activation and reduces vascular inflammation in endothelial cells. Biomed Pharmacother, 163, 114755.

Iwanaga, T., Kobayashi, D., Hirayama, M., Maeda, T. & Tamai, I. 2005. Involvement of uric acid transporter in increased renal clearance of the xanthine oxidase inhibitor oxypurinol induced by a uricosuric agent, benzbromarone. Drug Metab Dispos, 33, 1791–5.

Jankovsky, N., Caulier, A., Demagny, J., Guitton, C., Djordjevic, S., Lebon, D., Ouled-Haddou, H., Picard, V. & Garcon, L. 2021. Recent advances in the pathophysiology of PIEZO1-related hereditary xerocytosis. Am J Hematol, 96, 1017–1026.

Jansen, T. L., Reinders, M. K., Van Roon, E. N. & Brouwers, J. R. 2004. Benzbromarone withdrawn from the European market: another case of “absence of evidence is evidence of absence”? Clin Exp Rheumatol, 22, 651.

Kaestner, L., Egee, S., Connes, P., Bogdanova, A. Y. & Simmonds, M. J. 2025. Splenic filtration of red blood cells: Physics, chemistry, and biology need to go hand in hand. Proc Natl Acad Sci U S A, 122, e2405086121.

Kaestner, L. & Minetti, G. 2017. The potential of erythrocytes as cellular aging models. Cell Death Differ, 24, 1475–1477.

Kaestner, L., Schlenke, P., Von Lindern, M. & El Nemer, W. 2024. Translatable tool to quantitatively assess the quality of red blood cell units and tailored cultured red blood cells for transfusion. Proc Natl Acad Sci U S A, 121, e2318762121.

Karlsson, M., Zhang, C., Mear, L., Zhong, W., Digre, A., Katona, B., Sjostedt, E., Butler, L., Odeberg, J., Dusart, P., Edfors, F., Oksvold, P., Von Feilitzen, K., Zwahlen, M., Arif, M., Altay, O., Li, X., Ozcan, M., Mardinoglu, A., Fagerberg, L., Mulder, J., Luo, Y., Ponten, F., Uhlen, M. & Lindskog, C. 2021. A single-cell type transcriptomics map of human tissues. Sci Adv, 7.

Kowarz, E., Loscher, D. & Marschalek, R. 2015. Optimized Sleeping Beauty transposons rapidly generate stable transgenic cell lines. Biotechnol J, 10, 647–53.

Kuck, L., Kaestner, L., Egee, S., Lew, V. L. & Simmonds, M. J. 2025. Mechanotransduction mechanisms in human erythrocytes: Fundamental physiology and clinical significance. Channels (Austin), 19, 2556105.

Liang, P., Wan, Y. S., Shan, K. Z., Chou, R., Zhang, Y., Delahunty, M., Khandelwal, S., Francis, S. J., Arepally, G. M., Telen, M. J. & Yang, H. 2025. Targeting PIEZO1-TMEM16F Coupling to Mitigate Sickle Cell Disease Complications. Am J Hematol, 100, 2261–2275.

Liang, P., Zhang, Y., Wan, Y. C. S., Ma, S., Dong, P., Lowry, A. J., Francis, S. J., Khandelwal, S., Delahunty, M., Telen, M. J., Strouse, J. J., Arepally, G. M. & Yang, H. 2024. Deciphering and disrupting PIEZO1-TMEM16F interplay in hereditary xerocytosis. Blood, 143, 357–369.

Murciano, N., Rotordam, M. G., Becker, N., Ludlow, M. J., Parsonage, G., Darras, A., Kaestner, L., Beech, D. J., George, M., Fertig, N., Rapedius, M. & Bruggemann, A. 2023. A high-throughput electrophysiology assay to study the response of PIEZO1 to mechanical stimulation. J Gen Physiol, 155.

Nader, E., Conran, N., Leonardo, F. C., Hatem, A., Boisson, C., Carin, R., Renoux, C., Costa, F. F., Joly, P., Brito, P. L., Esperti, S., Bernard, J., Gauthier, A., Poutrel, S., Bertrand, Y., Garcia, C., Saad, S. T. O., Egee, S. & Connes, P. 2023. Piezo1 activation augments sickling propensity and the adhesive properties of sickle red blood cells in a calcium-dependent manner. Br J Haematol, 202, 657–668.

Obergrussberger, A., Haarmann, C., Rinke, I., Becker, N., Guinot, D., Brueggemann, A., Stoelzle-Feix, S., George, M. & Fertig, N. 2014. Automated Patch Clamp Analysis of nAChalpha7 and Na(V)1.7 Channels. Curr Protoc Pharmacol, 65, 11 13 1–48.

Pan, X., Wan, R., Wang, Y., Liu, S., He, Y., Deng, B., Luo, S., Chen, Y., Wen, L., Hong, T., Xu, H., Bian, Y., Xia, M. & Li, J. 2022. Inhibition of chemically and mechanically activated Piezo1 channels as a mechanism for ameliorating atherosclerosis with salvianolic acid B. Br J Pharmacol, 179, 3778–3814.

Parsonage, G., Cuthbertson, K., Endesh, N., Murciano, N., Hyman, A. J., Revill, C. H., Povstyan, O. V., Chuntharpursat-Bon, E., Debant, M., Ludlow, M. J., Futers, T. S., Lichtenstein, L., Kinsella, J. A., Bartoli, F., Rotordam, M. G., Becker, N., Bruggemann, A., Foster, R. & Beech, D. J. 2023. Improved PIEZO1 agonism through 4-benzoic acid modification of Yoda1. Br J Pharmacol, 180, 2039–2063.

Rapetti-Mauss, R., Picard, V., Guitton, C., Ghazal, K., Proulle, V., Badens, C., Soriani, O., Garcon, L. & Guizouarn, H. 2017. Red blood cell Gardos channel (KCNN4): the essential determinant of erythrocyte dehydration in hereditary xerocytosis. Haematologica, 102, e415–e418.

Rode, B., Shi, J., Endesh, N., Drinkhill, M. J., Webster, P. J., Lotteau, S. J., Bailey, M. A., Yuldasheva, N. Y., Ludlow, M. J., Cubbon, R. M., Li, J., Futers, T. S., Morley, L., Gaunt, H. J., Marszalek, K., Viswambharan, H., Cuthbertson, K., Baxter, P. D., Foster, R., Sukumar, P., Weightman, A., Calaghan, S. C., Wheatcroft, S. B., Kearney, M. T. & Beech, D. J. 2017. Piezo1 channels sense whole body physical activity to reset cardiovascular homeostasis and enhance performance. Nat Commun, 8, 350.

Romero, L. O., Bade, M., Elsherif, L., Williams, J. D., Kong, X., Adebiyi, A., Ataga, K. I., Ma, S., Cordero-Morales, J. F. & Vasquez, V. 2025. Enhanced PIEZO1 function contributes to the pathogenesis of sickle cell disease. Proc Natl Acad Sci U S A, 122, e2514863122.

Rosa, N., Ivanova, H., Wagner, L. E., 2ND, Kale, J., La Rovere, R., Welkenhuyzen, K., Louros, N., Karamanou, S., Shabardina, V., Lemmens, I., Vandermarliere, E., Hamada, K., Ando, H., Rousseau, F., Schymkowitz, J., Tavernier, J., Mikoshiba, K., Economou, A., Andrews, D. W., Parys, J. B., Yule, D. I. & Bultynck, G. 2022. Bcl-xL acts as an inhibitor of IP(3)R channels, thereby antagonizing Ca(2+)-driven apoptosis. Cell Death Differ, 29, 788–805.

Salmon, P. 2013. Generation of human cell lines using lentiviral-mediated genetic engineering. Methods Mol Biol, 945, 417–48.

Shan, Y., Guo, X., Zhang, M., Chen, M., Li, Y., Zhang, M. & Pei, D. 2025. Structure of human PIEZO1 and its slow-inactivating channelopathy mutants. Elife, 13.

Silvani, G., Kopecky, C., Romanazzo, S., Rodriguez, V., Das, A., Pandzic, E., Lock, J. G., Chaffer, C. L., Poole, K. & Kilian, K. A. 2025. Capillary constrictions prime cancer cell tumorigenicity through PIEZO1. Nat Commun, 16, 8160.

Smith, C. J., Zhang, Y., Koboldt, C. M., Muhammad, J., Zweifel, B. S., Shaffer, A., Talley, J. J., Masferrer, J. L., Seibert, K. & Isakson, P. C. 1998. Pharmacological analysis of cyclooxygenase-1 in inflammation. Proc Natl Acad Sci U S A, 95, 13313–8.

Syeda, R., Xu, J., Dubin, A. E., Coste, B., Mathur, J., Huynh, T., Matzen, J., Lao, J., Tully, D. C., Engels, I. H., Petrassi, H. M., Schumacher, A. M., Montal, M., Bandell, M. & Patapoutian, A. 2015. Chemical activation of the mechanotransduction channel Piezo1. Elife, 4.

Tripette, J., Alexy, T., Hardy-Dessources, M. D., Mougenel, D., Beltan, E., Chalabi, T., Chout, R., Etienne-Julan, M., Hue, O., Meiselman, H. J. & Connes, P. 2009. Red blood cell aggregation, aggregate strength and oxygen transport potential of blood are abnormal in both homozygous sickle cell anemia and sickle-hemoglobin C disease. Haematologica, 94, 1060–5.

Wang, Y., Chi, S., Guo, H., Li, G., Wang, L., Zhao, Q., Rao, Y., Zu, L., He, W. & Xiao, B. 2018. A lever-like transduction pathway for long-distance chemical- and mechano-gating of the mechanosensitive Piezo1 channel. Nat Commun, 9, 1300.

Woo, S. H., Ranade, S., Weyer, A. D., Dubin, A. E., Baba, Y., Qiu, Z., Petrus, M., Miyamoto, T., Reddy, K., Lumpkin, E. A., Stucky, C. L. & Patapoutian, A. 2014. Piezo2 is required for Merkel-cell mechanotransduction. Nature, 509, 622–6.

Xu, L., Li, T., Cao, Y., He, Y., Shao, Z., Liu, S., Wang, B., Su, A., Tian, H., Li, Y., Liang, G., Wang, C., Shyy, J., Xiong, Y., Chen, F., Yuan, J. X., Liu, J., Zhou, B., Wettschureck, N., Offermanns, S., Yan, Y., Yuan, Z. & Wang, S. 2025. PIEZO1 mediates periostin+ myofibroblast activation and pulmonary fibrosis in mice. J Clin Invest, 135.

Yaganoglu, S., Kalyviotis, K., Vagena-Pantoula, C., Julich, D., Gaub, B. M., Welling, M., Lopes, T., Lachowski, D., Tang, S. S., Del Rio Hernandez, A., Salem, V., Muller, D. J., Holley, S. A., Vermot, J., Shi, J., Helassa, N., Torok, K. & Pantazis, P. 2023. Highly specific and non-invasive imaging of Piezo1-dependent activity across scales using GenEPi. Nat Commun, 14, 4352.

Yeromin, A. V., Zhang, S. L., Jiang, W., Yu, Y., Safrina, O. & Cahalan, M. D. 2006. Molecular identification of the CRAC channel by altered ion selectivity in a mutant of Orai. Nature, 443, 226–9.

Zarychanski, R., Schulz, V. P., Houston, B. L., Maksimova, Y., Houston, D. S., Smith, B., Rinehart, J. & Gallagher, P. G. 2012. Mutations in the mechanotransduction protein PIEZO1 are associated with hereditary xerocytosis. Blood, 120, 1908–15.

Zhen, L., King, A. A., Xiao, Y., Chanock, S. J., Orkin, S. H. & Dinauer, M. C. 1993. Gene targeting of X chromosome-linked chronic granulomatous disease locus in a human myeloid leukemia cell line and rescue by expression of recombinant gp91phox. Proc Natl Acad Sci U S A, 90, 9832–6.

