## Supplementary figures and images for "Otenabant is a Selective Antagonist of Human PIEZO1"

### Supplementary figure 1

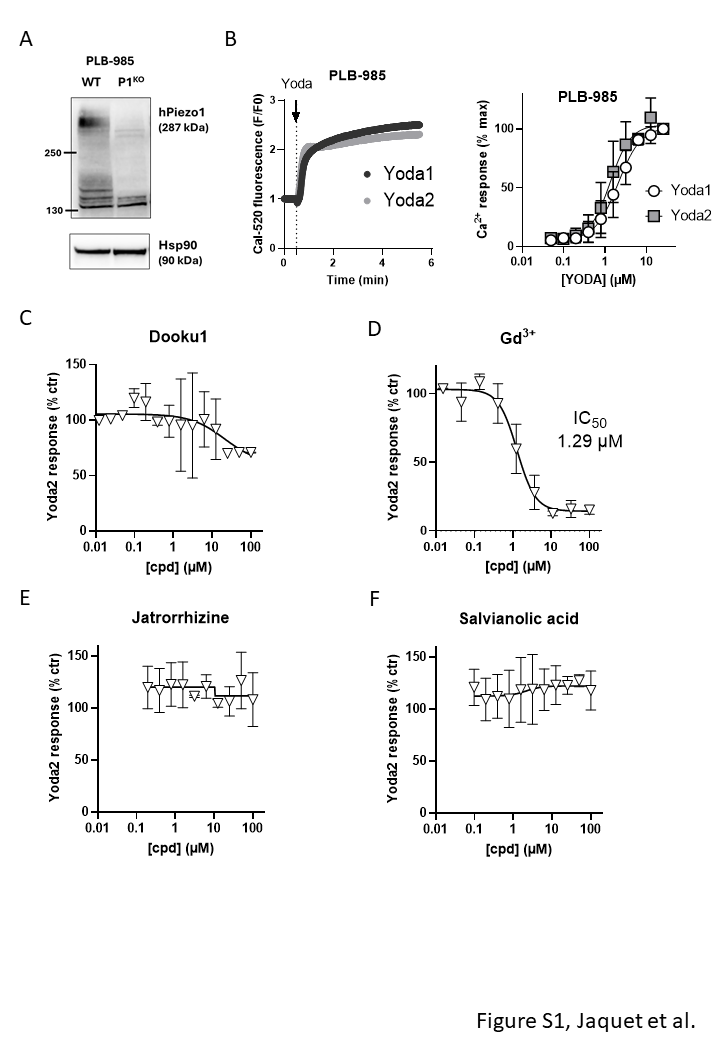

### Supplementary figure 2

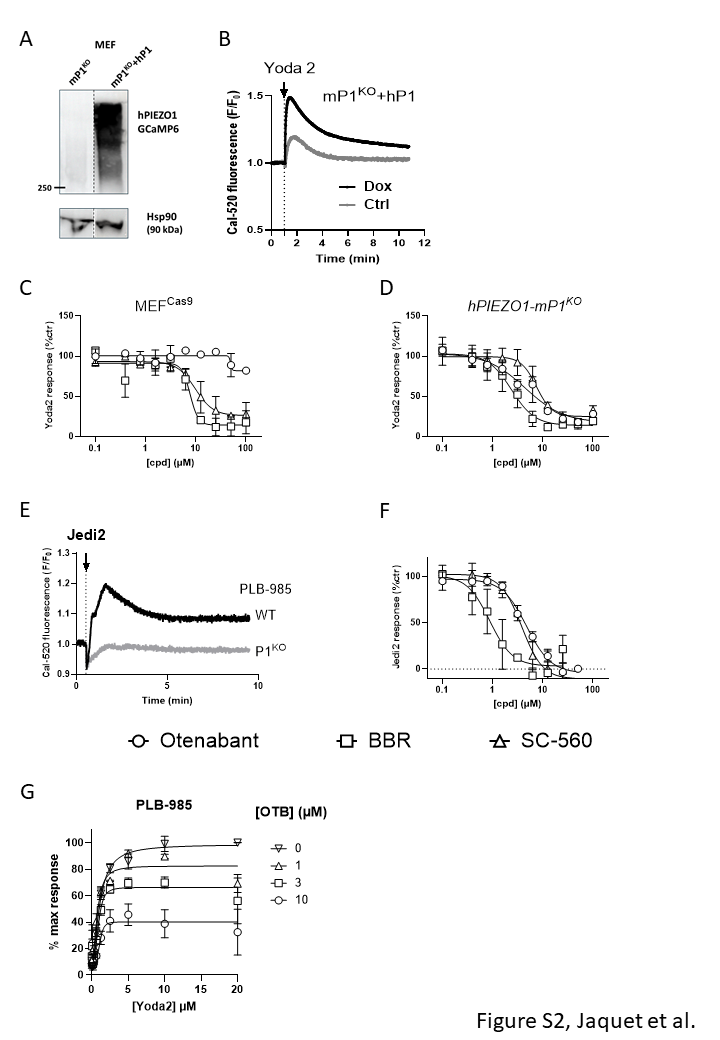

### Supplementary figure 3

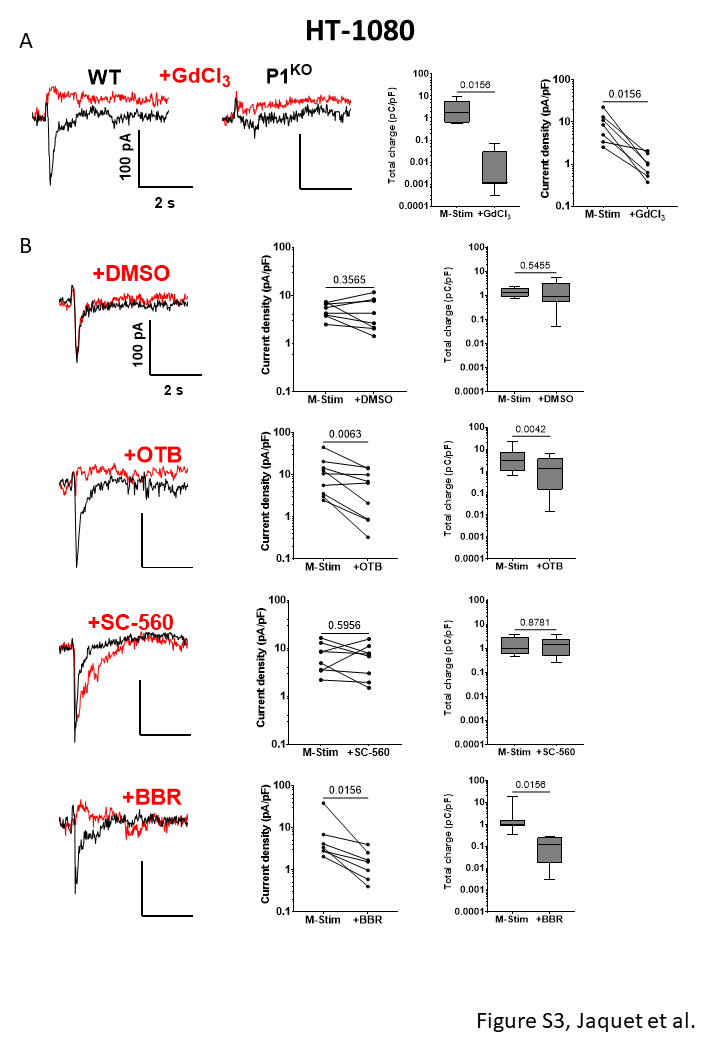

### Supplementary figure 5

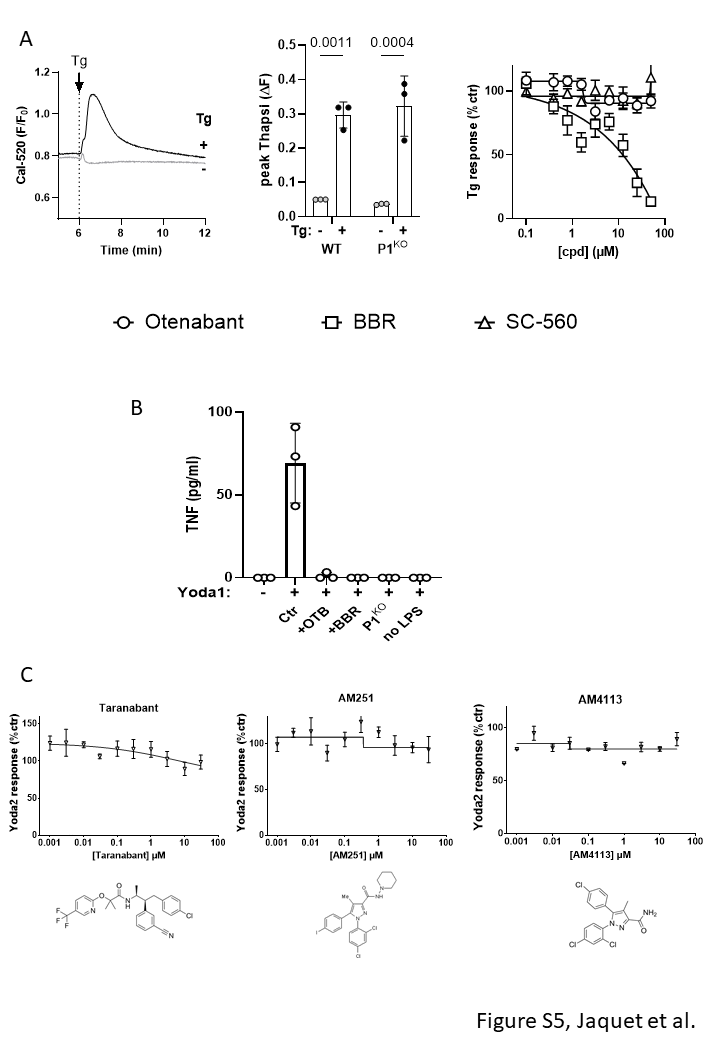

### Supplementary figure 6

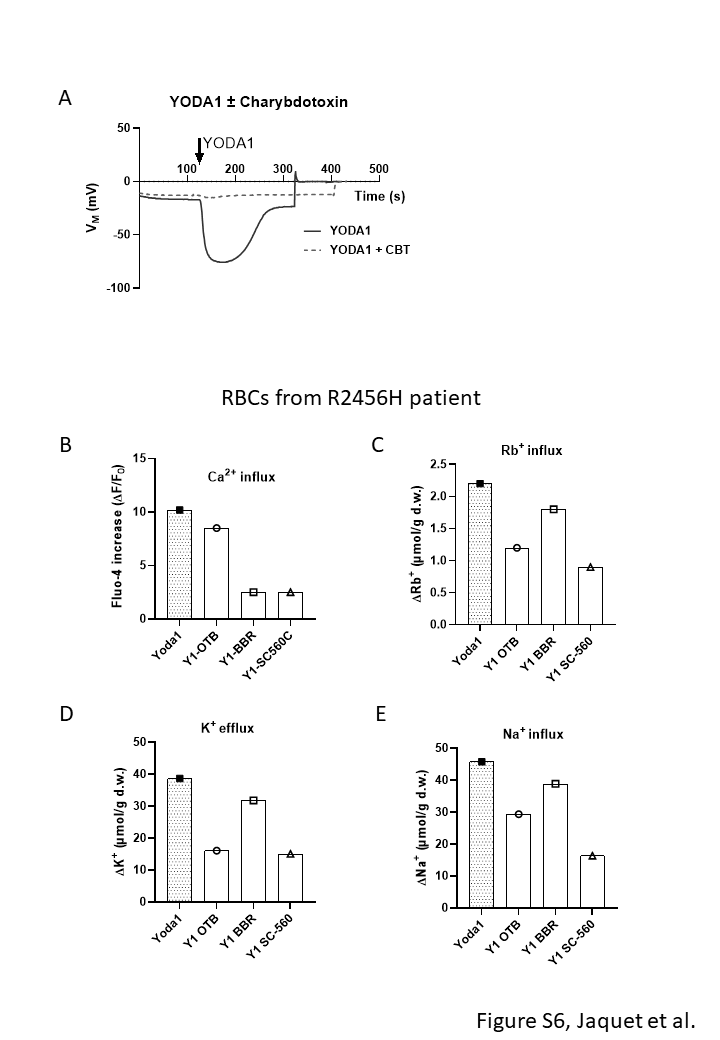
